# Extensive structural remodeling of the axonal arbors of parvalbumin basket cells during development

**DOI:** 10.1101/2021.03.03.433691

**Authors:** Kristina D. Micheva, Marianna Kiraly, Marc M. Perez, Daniel V. Madison

## Abstract

Parvalbumin-containing (PV+) basket cells are specialized cortical interneurons that regulate the activity of local neuronal circuits with high temporal precision and reliability. PV+ interneuron disfunction is associated with numerous psychiatric disorders, including schizophrenia and autism spectrum disorders. To understand how the PV+ interneuron connectivity underlying their functional properties is established during development, we used array tomography to map pairs of synaptically connected PV+ interneurons and postsynaptic neurons from the neocortex of mice of both sexes. We focused on the axon-myelin unit of the PV+ interneuron and quantified the number of synapses onto the postsynaptic neuron, length of connecting axonal paths, and their myelination at different time points between 2 weeks and 7 months of age. We find that myelination of the proximal axon occurs very rapidly during the third postnatal week and precedes a massive synapse pruning which takes place in the 4th postnatal week leading to about three-fold reduction of synaptic contacts made by the PV+ interneuron on its postsynaptic partner. Autapses, the synapses that PV+ interneurons form on themselves, however, are not subjected to pruning. Axon reorganizations continue beyond postnatal month 2, with the postsynaptic targets of PV+ interneurons gradually shifting to more proximal locations, and the length of axonal paths and their myelin becoming conspicuously uniform per connection. These continued microcircuit refinements likely provide the structural substrate for the robust inhibitory effects and fine temporal precision of PV+ basket cells.

**Significance statement:** The axon of adult parvalbumin-containing (PV+) interneurons is highly specialized for fast and reliable neurotransmission. It is myelinated and forms synapses mostly onto the cell bodies and proximal dendrites of postsynaptic neurons for maximal impact. In this study we follow the development of the PV+ interneuron axon, its myelination and synapse formation, revealing a rapid sequence of axonal reorganization, myelination of the PV+ interneuron proximal axon, and subsequent pruning of almost two-thirds of the synapses in an individual connection. This is followed by a prolonged period of axon refinement and additional myelination leading to a remarkable precision of connections in the adult mouse cortex, consistent with the temporal precision and fidelity of PV+ interneuron action.

## Introduction

Parvalbumin positive basket cells are the most abundant type of inhibitory interneurons in neocortex. Found in all cortical layers, except layer 1, and in all cortical areas (Rudy et al., 2011), they form synapses onto hundreds of neighboring pyramidal neurons and interneurons (Packer et al., 2011) targeting preferentially their cell bodies and proximal dendrites (Somogyi et al., 1983; Tamás et al., 1997a; Kawaguchi and Kubota, 1998). This connectivity underlies the many important roles of PV+ basket cells, such as controlling cortical rhythmic activity (Cardin et al., 2009; Buzsáki and Wang, 2012; Cardin 2018; Chariker et al., 2018), regulating the excitatory/inhibitory balance (Xue et al., 2014; Ferguson and Gao, 2018) and controlling spike timing of pyramidal neurons (Pouille and Scanziani, 2001; Wehr and Zador, 2003). The initial connectivity of PV+ interneurons is established rapidly during the early postnatal ages, followed by circuit refinement that underlies learning and plasticity and continues through adult life. During the first postnatal week in mouse neocortex, the majority of PV+ interneurons migrate to their final laminar destination and form synapses with neighboring neurons (Lim et al., 2018). Then, towards the end of this first week, many PV+ interneurons begin to die off, and by P10 as many as 40% of the PV+ interneurons, thought to be the ones that did not successfully integrate in the developing neuronal circuits (Southwell et al., 2012, Wong et al., 2018), undergo apoptosis. Around the same time, profound changes are seen in the expression of numerous genes in PV+ interneurons, including genes involved in synaptic function (Okaty et al., 2009). PV+ expression appears around P12-P14 (del Río et al., 1994). Electrophysiological properties of PV+ interneurons also change dramatically, including the loss of spike-frequency adaptation between P10 and P15 and large increase in the maximum firing rate (Okaty et al., 2009; Miyamae et al., 2017). Thus, within the span of less than 2 weeks, neocortical PV+ basket cells arrive at their final destination, establish synaptic connections and integrate into the developing neocortical circuit or die off.

While the genetic programs and molecular mechanisms of this initial development of neocortical PV+ interneurons have been extensively characterized (Lim et al., 2018), much less is known about the synaptic connectivity of the surviving PV+ interneurons at this stage and the subsequent reorganization of their axonal arbors during circuit refinement. During the second half of the first postnatal month, the number of the perisomatic synapses that PV+ interneurons form around nearby pyramidal neurons gradually increases (Chattopadhyaya et al., 2004). This is the net result of on-going new synapse formation and synapse elimination, in an activity-dependent manner (Chattopadhyaya et al., 2004, Wu et al., 2012). At the same time, as these interneurons mature, their proximal axonal arbor becomes myelinated in a characteristic patchy manner (Zonouzi et al., 2019). How do these events unfold relative to one another? When does the myelination take place? How do myelination and synaptic formation/pruning reshape the axonal arbor? These are the questions that our current study addresses. Using array tomography we map the connectivity in pairs of synaptically connected PV+ interneurons and postsynaptic neurons and quantify the number of synapses, length of connecting axonal paths and their myelination at different time points between 2 weeks of age and 7 months. We find that myelination of the proximal axon occurs very rapidly during the third postnatal week. It precedes the massive synapse pruning which takes place in the 4^th^ postnatal week and results in about three-fold reduction of synaptic contacts made by the PV+ interneuron on its postsynaptic partner. In contrast, the number of autapses, the synapses that the PV+ interneuron makes on itself, remain constant throughout this time. Our study reveals the extensive reorganization of the PV+ interneuron axonal arbor and synapses resulting in the remarkably precise adult connectivity pattern characterized by synapses proximal to the postsynaptic cell body and a uniform length of axonal paths to these synapses.

## Materials and Methods

### Animals

Young mice C57BL/6 of both sexes, ages P14 and P22, were used for this study. All procedures related to the care and treatment of animals were carried out in accordance with the Guide for the Care and Use of Laboratory Animals of the National Institutes of Health and approved by the Administrative Panel on Laboratory Animal Care at Stanford University. To study the developmental sequence, we also analyzed data from additional 14 pairs at ages spanning up to 7 months (1 pair at P22, 4 pairs at 1 month, 5 pairs at 2 months and 4 pairs at 7 months) used in a previous study (Micheva et al., 2021).

### Preparation of Acute Neocortical Slices

Brain slices were prepared exactly following the protocol of Ting et al., 2018. Briefly, mouse brains were hemisected, embedded in 2% agarose and cut to a thickness of 300 μm using a Compresstome vibrating microtome (Precisionary Instruments). Slices were stored submerged for at least one hour at room temperature in a Model 4 Brain Slice Keeper (AutoMate Scientific). During this incubation period, slices were subjected to the Wisteria floribunda lectin staining procedure below. For recording, slices were transferred to a submerged/superfusing slice chamber with a glass coverslip bottom (Warner Instruments) on the stage of an upright microscope.

### PV+ Interneuron identification

To identify PV+ interneurons in living tissue, the perineuronal nets that selectively surround these neurons were stained in the live slices using fluorescein labeled Wisteria floribunda lectin (Vector Labs FL-1351), following the protocol of Hoppenrath et al., 2016. Briefly, immediately after sectioning on the Compresstome, slices were transferred into a small volume of holding buffer (in mM: 92 NaCl, 2.5 KCl, 1.25 NaH2PO4, 30 NaHCO3, 20 HEPES, 25 glucose, 2 thiourea, 5 Na-ascorbate, 3 Na-pyruvate, 2 CaCl2·2H2O, and 2 MgSO4·7H2O, pH to 7.3–7.4) containing fluorescein labeled Wisteria floribunda lectin (20 μg/ml) and maintained at room temperature under a 95% O_2_, 5 % CO_2_ atmosphere for 1 hour. Labeled perineuronal nets were detected in the live slices using epiflourescence (Nikon GFP filter cube). The identity of the neurons labeled by the Wisteria floribunda lectin was previously confirmed by us using subsequent immunofluorescent labeling with anti-Parvalbumin antibody (Micheva et al., 2021).

### Electrophysiology

We performed simultaneous whole-cell recordings from two different neurons from the mouse medial temporal lobe (perirhinal and entorhinal cortex). The presynaptic PV+ basket cell was identified by Wisteria floribunda labeling of its perineuronal net and a nearby postsynaptic pyramidal neuron was chosen by its shape. Neurons at a 30 to 50 μm depth were targeted, because healthy neurons were typically located below 30 μm of depth within the slice (Ting et al., 2018), and Wisteria floribunda labeling was usually limited within the surface 50 μm. The presynaptic interneuron was recorded in current clamp, and the postsynaptic neuron in single-electrode continuous voltage clamp (Molecular Devices Multiclamp 700A). Single action potentials were elicited from the presynaptic interneuron by the injection of positive current through the recording electrode, with the amplitude and duration of the current pulse adjusted to produce a single action potential. Data were sampled at 10 KHz and low pass filtered between 3 and 10 KHz. The postsynaptic recording was examined for time-locked postsynaptic responses to that presynaptic action potential. Typically, at least 25 action potentials/IPSC trials were recorded per cell. We measured the amplitude of the evoked postsynaptic current and the latency of that current, relative in time to the peak of the presynaptic action potential. The detailed methods, including recording configurations, solutions, electrodes, etc. were as in (Pavlidis and Madison, 1999, Montgomery et al., 2001), except that the postsynaptic electrode internal solution consisted of (in mM): 120 KCl, 40 HEPES, 2 Mg-ATP, 0.317 Na-GTP, and 10.7 MgCl_2_. Besides making the IPSCs larger and easier to detect, use of high chloride solution also reversed the usual polarity of inhibitory synaptic currents to inward. To mark the neurons for subsequent AT analysis, the presynaptic interneuron internal electrode solution included 0.1% Alexa Fluor 594 hydrazide (ThermoFisher Scientific A-10438) and 0.5% Neurobiotin (Vector Laboratories SP-1120-50), and the postsynaptic neuron internal electrode solution contained 0.2% Lucifer Yellow CH, potassium salt (ThermoFisher Scientific L1177).

### Slice fixation and embedding in resin

After the electrophysiological recordings, the slice was placed in a fixative containing 2% formaldehyde (Electron Microscopy Sciences 157-8) and 2 % glutaraldehyde (Electron Microscopy Sciences 16220) in PBS, first at room temperature for 1 hour, and then at 4°C for 24h. Slices were subsequently washed in PBS, and a smaller area (approximately 1 by 2 mm) around the labeled neurons was dissected out and dehydrated serially in washes of 50%, 70%, 70% ethanol, at 4°C, 10 min for each step. The dehydrated tissue was then infiltrated with LR White resin, hard grade (Electron Microscopy Sciences 14383), first with a mixture of 70% ethanol and LR White (1:3) and then in three changes of 100% LR White, 10 min at room temperature each step. The slice was left in unpolymerized LR White resin overnight at 4°C, then transferred to a gelatin capsule size “00” (Electron Microscopy Sciences 70100) and polymerized for 24h at 55°C. The polymerized blocks with tissue were stored in the dark at room temperature.

### Sectioning

The blocks were trimmed around the tissue to the shape of a trapezoid approximately 1.5 mm wide and 0.5 mm high, and glue (Weldwood Contact Cement diluted with xylene) was applied with a thin paint brush to the leading and trailing edges of the block pyramid. The embedded plastic block was cut on an ultramicrotome (Leica Ultracut EM UC6) into 100-nm-thick serial sections, which were mounted on gelatin-coated coverslips. Typically, 1,000 to 1,200 serial sections containing the fixed tissue slice were obtained from each block.

### Immunofluorescent array tomography

Coverslips with sections containing the filled neurons were identified by visual inspection using a 10x objective under the fluorescence microscope, and were processed for standard indirect immunofluorescence, as described in Micheva et al., 2010. Antibodies were obtained from commercial sources and are listed in Table 1. The sections were pretreated with sodium borohydride [1% in Tris-buffered saline (TBS), pH 7.6 for 3 min] to reduce non-specific staining and autofluorescence. After a 20 min wash with TBS, the sections were incubated in 50 mM glycine in TBS for 5 min, followed by blocking solution (0.05% Tween 20 and 0.1% BSA in TBS) for 5 min. The primary antibodies were diluted in blocking solution as specified in Table 1 and were applied overnight at 4°C. After a 15-min wash in TBS, the sections were incubated with Alexa Fluor dye-conjugated secondary antibodies, highly cross-adsorbed (ThermoFisher Scientific), diluted 1:150 in blocking solution for 30 min at room temperature. Neurobiotin was detected using Streptavidin-Alexa 594 (ThermoFisher Scientific S11227), diluted at 1:200, and applied together with the secondary antibodies. Finally, sections were washed with TBS for 15 min, rinsed with distilled water, and mounted on glass slides using SlowFade Diamond Antifade Mountant with DAPI (ThermoFisher Scientific S36964). After the sections were imaged, the antibodies were eluted using a solution of 0.2 M NaOH and 0.02% SDS, and new antibodies were reapplied on some of the coverslips, as needed. The first round of staining always included anti-LY and Streptavidin-Alexa 594 to visualize the filled neurons and anti-MBP to detect myelin.

**Table 1.** Antibodies.

| Antigen | Host | Antibody Source | Dilution | RRID |
| --- | --- | --- | --- | --- |
| Lucifer Yellow | rabbit | ThermoFisher Scientific A-5750 | 1:200 | AB_2536190 |
| Myelin basic protein | chicken | AVES, MBP | 1:100 | AB_2313550 |
| Synapsin 1/2 | chicken | Synaptic Systems 106 006 | 1:200 | AB_2622240 |
| Gephyrin | mouse | NeuroMab clone L106/83, 75-443 | 1:100 | AB_2636851 |
| Pan-QKI | mouse | NeuroMab clone N147/6, 73-168 | 1:2 | AB_10671658 |

### Imaging

The immunostained ribbons of sections were imaged on an automated epifluorescence microscope (Zeiss AxioImager Z1) with AxioVision software (Zeiss) using a 63x Plan-Apochromat 1.4 NA oil objective. To define the position list for the automated imaging, a custom Python-based graphical user interface, MosaicPlanner (obtained from https://code.google.com/archive/p/smithlabsoftware/), was used to automatically find corresponding locations across the serial sections. On average, 582 sections per neuron pair were imaged (ranging between 375 to 826 sections), resulting in volumes of approximately 140 μm by 400 μm by 58 μm centered on the filled neurons. Images from different imaging sessions were registered using a DAPI stain present in the mounting medium. The images from the serial sections were also aligned using the DAPI signal. Both image registration and alignment were performed with the MultiStackReg plugin in FIJI (Schindelin et al., 2012).

### Synapse detection and axon tracing

A putative synapse between the two filled neurons was identified as a direct apposition between an enlargement in the presynaptic axon and the postsynaptic neuron, which was present on two or more consecutive sections. Twenty-two putative synapses from three different P14 pairs were subsequently immunostained with synaptic markers to verify whether these criteria were adequate to identify synapses. The expression of synaptic markers was lower at this age compared to older animals (Micheva et al., 2021), and gephyrin immunoreactivity in particular was ambiguous to assess in many cases (12 out of 22 contacts) and was therefore not used for this analysis. Out of 22 contacts at this age, 20 were synapsin immunopositive. At P22, out of 16 contacts, all were synapsin immunopositive. For brevity, in the following sections the putative synaptic contacts in the imaged pairs are referred to as synapses. Using FIJI, axons were traced back from the synapses to the neuronal cell body. Each connection was traced independently by two experts (MMP and KDM) and axonal path trajectories were saved as a set of ROIs. Myelinated internodes were identified using MBP immunofluorescence, which is a reliable measure of myelin as previously verified using correlative immunofluorescence – SEM (Micheva et al., 2016). Axon diameter was estimated as the average of three measurements (beginning, middle and end of each internode), using the measure function in FIJI. Axon initial segment diameter was also calculated as the average of three measurements along its length. Altogether, the following morphological measurements were obtained: total axonal distance to each synapse, length and diameter of axon initial segment, length, diameter and number of myelinated internodes, length of nodes of Ranvier, distance of the postsynaptic target from the postsynaptic cell body. The results for each pair were visualized as maps of the PV+ interneuron axonal arbors (axonograms) and maps of the postsynaptic neuron dendritic arbors with location of the PV+ synapses (dendrograms, graphical representation inspired by Aliaga Maraver et al., 2018). Because tissue dehydration and embedding with our protocol result in approximately 23% linear shrinkage, these measurements were adjusted accordingly (Busse and Smith, 2013).

### Statistics

Measurements are presented as average ± standard deviation. For comparison between groups we used two-tailed unpaired t-test. Linear correlations between 2 variables were done using Pearson’s correlation coefficient. To prepare the boxplots, the web application BoxPlotR was used (http://boxplot.tyerslab.com; Spitzer et al. 2014).

## Results

To analyze the developmental changes in connectivity pattern and axonal myelination of PV+ interneurons we recorded from seventeen synaptically connected pairs of neurons from 11 mice (14 pairs from 9 mice at P14, and 3 pairs from 2 mice at P22). The presynaptic neuron was always a PV+ interneuron identified by the fluorescent Wisteria floribunda stain of its perineuronal net, and the postsynaptic neuron was an adjacent pyramidal neuron. Six of these pairs (4 at P14 and 2 at P22) that had bright and complete neuronal fills were further processed for immunofluorescent array tomography to find the synapses in each connection and map the connecting axonal arbor. The results were compared with data from additional 14 pairs at ages spanning up to 7 months (1 pair at P22, 4 pairs at 1 month, 5 pairs at 2 months and 4 pairs at 7 months) that were used in a previous study (Micheva et al., 2021).

### Profuse synaptic connectivity of PV+ interneurons at P14, but scarce myelination of their axons

At the youngest age examined, P14, the interneuronal axon branched profusely and formed numerous axonal varicosities (Figures 1 and 2). These varicosities were densely distributed on 3^rd^ and higher order branches at an average of 3.7 ± 0.6 varicosities per 10 μm of axonal length (average ± standard deviation) and the great majority of them (96%) were immunopositive for synapsin. In individual neuronal pairs, the PV+ interneuron formed on average 19 ± 6 synapses onto the postsynaptic pyramidal neuron. This is about twice as many synapses as found in older pairs (9 ± 5 synapses in pairs between 1 to 7 months; Micheva et al., 2021). Similarly to adult PV+ interneurons, the synapses were preferentially distributed close to the postsynaptic soma, with an average distance of 32 μm ± 24 μm from the soma, although a few synapses were seen as far away as 100 μm. In addition to the soma, synapses were formed both on the basal and apical dendrites, and, in several occasions (3 out of 76), on the axon initial segment. In each neuronal pair, the PV+ interneuron synapsed onto multiple postsynaptic domains (5 ± 2) of the postsynaptic neuron via multiple axonal paths (8 ± 2). A neuronal domain was defined as either the cell body, or a primary dendrite with its branches. In cases were the primary dendrite branched into two large branches close to the cell body (within 20 μm), these two branches were considered as separate domains. Axonal paths were considered as distinct if they were separated by 2 or more branching points.

**Figure 1.**
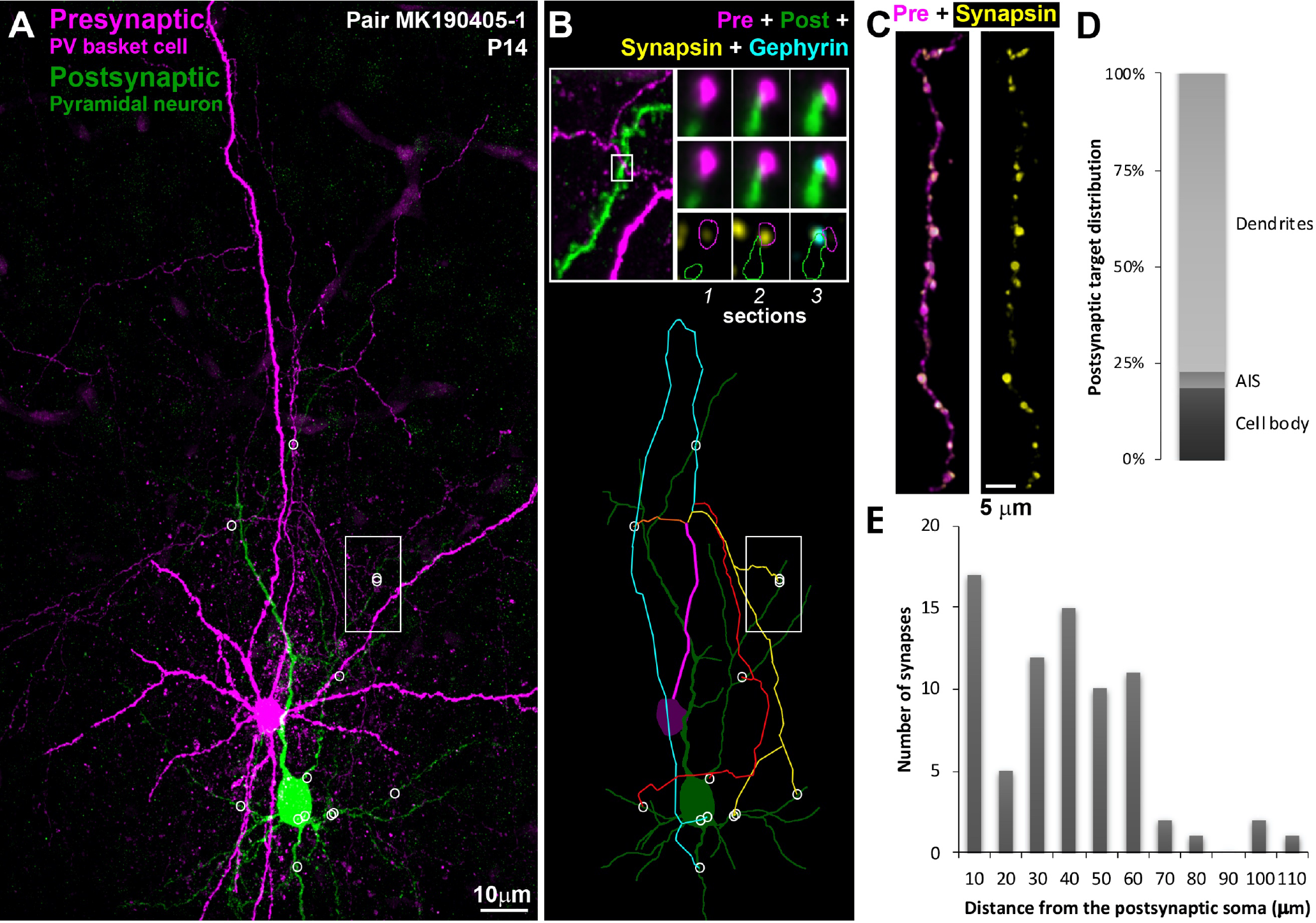
Synaptically connected PV+ interneuron – pyramidal neuron pairs at P14. **A.** MAX projection from pair MK190405-1, 480 serial ultrathin sections (100 nm each). Following ultrathin sectioning, the neurobiotin-filled presynaptic PV+ interneuron was labeled with Streptavidin Alexa 594 (magenta) and the Lucifer Yellow-filled pyramidal neuron with anti-Lucifer Yellow antibody (green). Locations of synapses are marked with white circles. The white rectangle identifies the area shown at higher magnification in B, top. **B.** Top Left, The marked area from A showing the location of a synapse between the interneuronal axon and an apical dendritic branch of the pyramidal neuron. Top, Right, Three serial sections through the synapse identified to the left. The presynaptic bouton is immunoreactive for synapsin (yellow) on sections 1 and 2, and gephyrin immunofluorescence (cyan) is observed at the postsynaptic side on section 3. Bottom, Map of the connection, with synapses represented by white circles. There are 4 main axonal paths (represented by different colors) in this connection. **C.** A stretch of the PV+ interneuron axon immunolabeled with synapsin (yellow). **D.** Distribution of postsynaptic targets at P14. **E.** Frequency distribution of the distances of each synapse from the postsynaptic cell body in the P14 connections (n=4).

**Figure 2.**
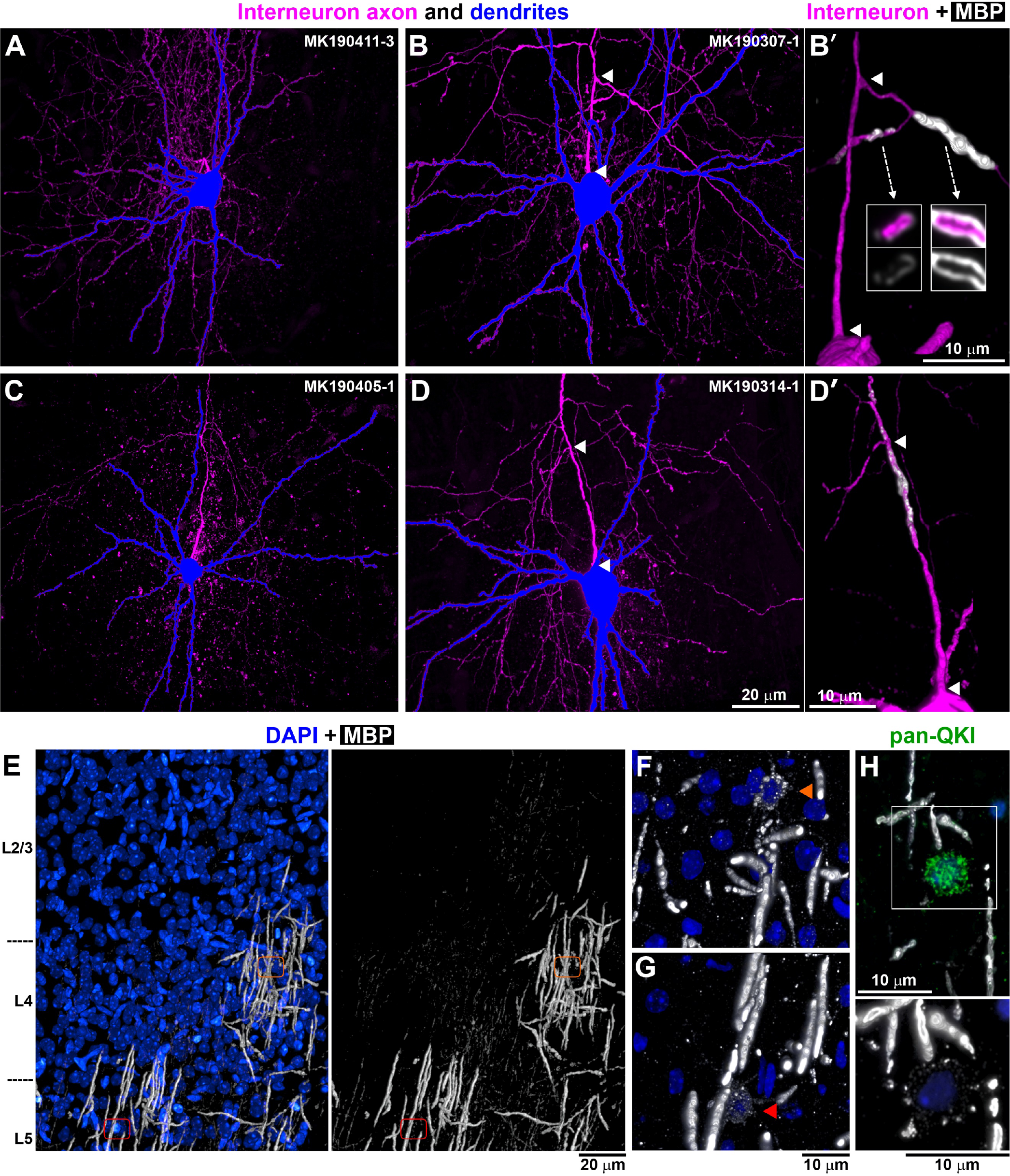
Axonal branching and myelination at P14. **A -D.** PV+ interneurons from P14 neuronal pairs, with the soma and dendrites in blue and axon in magenta. MAX projections through all sections from the pairs. **B’** and **D’** show volume reconstruction of the axon initial segment of interneurons B (103 sections) and D respectively (136 sections). Arrowheads mark the corresponding beginning and end of the first axonal segment from the soma to the first branching point. In B’ there are two tertiary axonal branches that are myelinated, the one going left is incompletely myelinated as seen by the uneven and weak MBP immunolabel. Arrows point to single sections through the axonal branches. No myelination is observed before the first branching point. In D’ there is incomplete myelination before the first branching point as well as after that. No complete myelin was seen in this PV+ interneuron. **E.** MBP immunostaining (white) – volume reconstruction from 529 sections. DAPI labeled nuclei are shown in blue. Areas with complete, incomplete and no MBP immunolabel are seen at this age. Orange and red outlines mark the locations of two different myelinating oligodendrocytes shown at higher magnification in F and G. **F, G**. Myelinating oligodendrocytes characterized by weak MBP immunolabel within their cell bodies and proximal cell processes. Volume reconstruction from 36 sections (F) and 73 sections (G). **H.** Weakly MBP-immunofluorescent oligodendrocytes (bottom, 13 sections volume reconstruction) co-labeled with an antibody against panQKI, an oligodendrocytic marker, (top, lower magnification).

In contrast to the abundant synapses, there was only minimal myelination of PV+ axons at P14 (Figure 2). Myelin was detected using immunofluorescence for myelin basic protein (MBP) that is highly enriched in myelin and reliably stains myelinated sheaths (Micheva et al., 2016). While in adult animals the MBP immunolabel precisely outlines myelinated stretches of axons, the internodes, at P14 a discontinuous and weak MBP labeling was observed around some axons suggesting myelination in progress, and we refer to it here as incomplete myelin. Examples of complete (adult-like) and incomplete myelin are shown in Figure 2B’,D’. Out of the four P14 pairs, one PV+ interneuron had no myelin on its axon, 2 interneurons had only incomplete myelin on a few axonal stretches, and one interneuron had both complete and incomplete myelin. The process of myelination does not appear to be occurring in the order of axonal branches, for example in pair MK190307, some third, fourth and fifth order axonal branches were myelinated, but no first or second order branches. This may be due to asynchronous maturation of the myelinating oligodendrocytes in upper cortical layers at this age. While at P14 myelin formation is observed in a rather uniform fashion in deeper layer 5 and in layer 6, MBP immunolabel in layers 2/3 and 4 revealed a mosaic distribution of regions with varying levels of myelination (complete myelin sheaths, incomplete myelin sheaths and no myelin sheaths, Figure 2E). Because oligodendrocytes, similarly to other types of glia, show tiling such that the territories of different cells do not overlap (Hughes et al., 2013), this distinct MBP pattern likely corresponds to the tiled distribution of oligodendrocytes at slightly different stages of maturation. Indeed, individual oligodendrocytes are known to myelinate axons preferentially within 50μm of their cell bodies (Murtie et al., 2007), and this roughly corresponds to the size of the regions with different myelination patterns that we observe (Figure 2E). Accordingly, within each tile with complete myelin, a myelinating oligodendrocyte could be discerned by its weak MBP immunolabel. Such weakly MBP positive glial cells co-labeled with the oligodendrocytic marker QKI (Figure 2H). Thus, the timing of myelination of different axonal branches may depend on the maturation level of the oligodendrocytes in the area, and not on their position in the axonal arbor.

### Extensive myelination occurs in the 3^rd^ postnatal week followed by substantial synapse pruning in the 4^th^ postnatal week

The above analysis reveals that compared to older mice (1 month of age and older, Micheva et al., 2021), the PV+ interneurons at P14 have little or no myelin, but abundant synaptic connections. When do the axonal arbors of these interneurons change and how do myelination and synaptic pruning unfold in time? To better understand the sequence of these events we analyzed 3 PV+ interneuron pairs from P22 mice (Figures 3, 3-1, 3-2 and 4) and then compared the P14 and P22 mice from this study with older mice from Micheva et al., 2021. Compared to P14 mice (19 ± 6 synapses), the number of synapses in a connection was essentially the same at P22 (16 ± 3), then dropped almost three-fold to 6 ± 2 at 1 month. Interestingly, the number of synapses subsequently almost doubled again to 10 ± 6 at 2 months and 10 ± 3 at 7 months, but the change was not statistically significant because of the large interpair variability at the older ages (p = 0.15 for 1 vs. 2 months, and p = 0.08 for 1 vs. 7 months; Figure 4A). To further validate these findings we also analyzed the electrophysiological data from all neuronal pairs (only a fraction of these was used for AT analysis as stated in the Methods). The amplitude of the presynaptic action potential-evoked inhibitory postsynaptic current (IPSC) correlates well with the number of synapses in a pair (Micheva et al., 2021), and in support of our AT results we found that the IPSC amplitude showed a developmental pattern similar to the number of synapses, with a statistically significant drop between P22 and 1 month of age (Figure 4B). On the other hand, the number of synaptic varicosities per axonal length showed a slight decrease between P14 and P22, after which it remained constant throughout development (Figure 4C), suggesting that the changes in synapse numbers are likely due to changes in axonal length.

**Figure 3.**
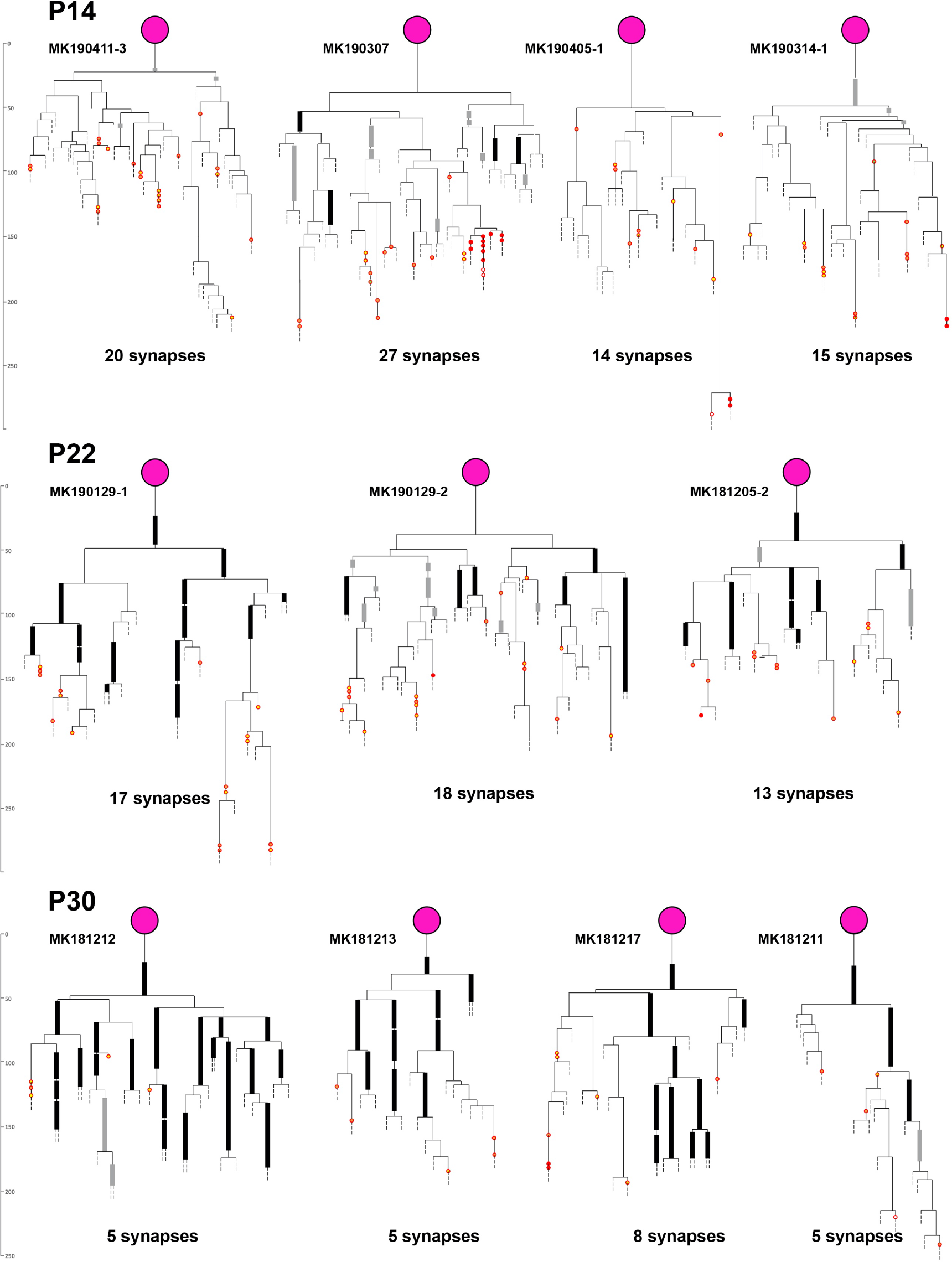
Axonograms of P14, P22 and P30 PV+ interneurons. Myelinated internodes are represented by thick black lines and incomplete myelination is shown with thick grey lines. Synapses are shown with red circles (full red – on soma, orange with red – on dendrite, yellow with red – on spine, white with red – on AIS). Information on the neuronal pairs is shown in Figure 3-1 and detailed axonograms and dendrograms of P14 and P22 pairs are shown in Figure 3-2.

**Figure 4.**
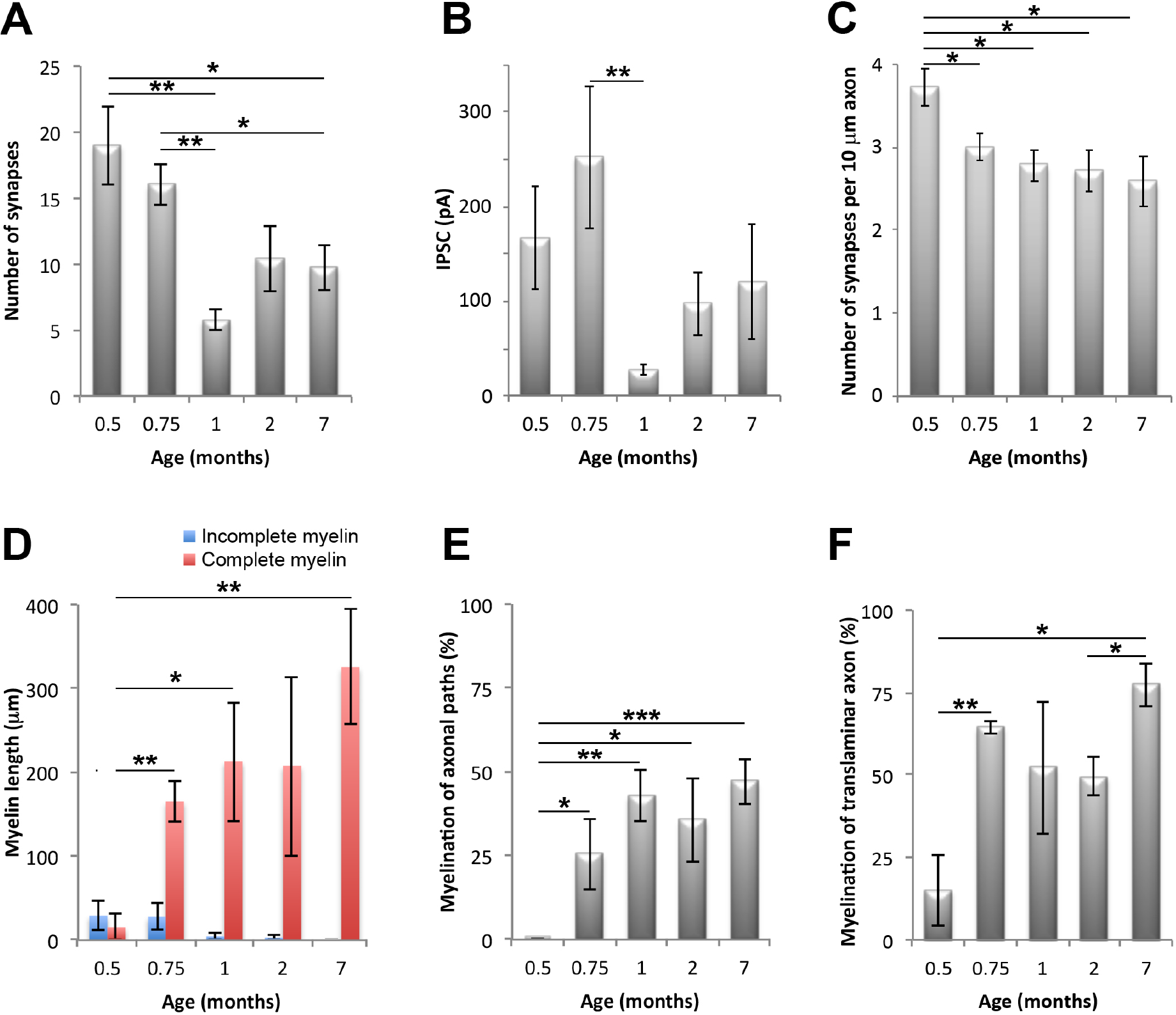
Changes in synapse numbers and in myelination of axonal paths of PV+ interneuron connections during development. **A.** Developmental changes in the number of synapses per PV+ interneuron connection. **B.** Changes in the amplitude of IPSC (n = 14 at P14, 4 at P22, 8 at 1 mo, 11 at 2 mo and 8 at 7 mo). **C.** Changes in the number of synaptic varicosities per axonal length. **D.** Changes in the total length of myelinated axon in the proximal 150 μm of the axonal arbor. **E.** Changes in the myelination of axonal paths in the connections. **F.** Changes in the myelination of the translaminar axonal branch. For all plots statistical significance is **\***P ≤ 0.05, **\*\***P ≤ 0.01. **\*\*\***P ≤ 0.001.

The changes in myelination occurred earlier than the initial decrease in synapse numbers, with a more than a ten-fold increase in the extent of myelin in the proximal 150 μm axonal arbor between P14 (15 ± 31 μm myelinated axon length per neuron) and P22 (165 ± 42 μm) and no statistically significant change at 1 month (213 ± 143 μm, p = 0.15). The incomplete myelin was still observed at P22, and then only occasionally at 1 and 2 months of age (Figure 4D).

### Proximal myelination is mostly complete within the first postnatal month, but myelination of the translaminar axonal branch continues much later in development

The percent myelination of the axonal paths in the connections jumped ten-fold between P14 (0.23 ± 0.03%) and P22 (25 ± 2%), further increasing at 1 month to 43 ± 2% (not statistically significant, p = 0.23) (Figure 4E). The above analysis was done by comparing the average values of the axonal paths per neuron, however, if comparing individual axonal paths at different ages, the changes between P22 and 1 month of age were significant (p = 0.03 for axonal paths in the connections). No significant changes were observed after 1 month. This was somewhat surprising considering that myelination is known to extend over a much longer period during development (Hill et al., 2018; Hughes et al., 2018). However, the pairs included in this analysis are only between neurons that are in the same cortical layer and in close proximity (40 μm or less between the two cell bodies geometric centers). In addition to the axonal arbor surrounding the soma, most of the analyzed interneurons also had a thick apical axonal branch extending upwards, likely synapsing onto neurons in upper layers (n=2 at P14 and n=3 at P22, 1 month, 2 months and 7 months each). The myelination of this apical axon continued at much later ages with significant differences observed between 2 months when it was 49% myelinated and 7 months with 77% myelination (p = 0.03, Figure 4F). Beyond the unmyelinated axon initial segment, which is included in the above estimates, the apical axon was almost completely myelinated at 7 months (92%), compared to 61% myelinated at 2 months (p = 0.02) Thus, while axonal paths to adjacent neurons in the same layer become rapidly myelinated in the 3^rd^ postnatal week, reaching adult levels by 1 month, myelination of the longer translaminar axonal branch continues much later, after 2 months.

### Following synapse pruning, adult connections consist of more proximally located synapses

Between P22 and 1 month there is an almost 3-fold decrease in the number of synapses between a PV+ interneuron and its postsynaptic target. How does this affect the topology of the connections? We first analyzed the developmental changes in the distribution of the postsynaptic targets. Initially, between P14 and P22 there was an increase in the average distance from the synapses to the postsynaptic soma; this may be due to the continuing growth of cortical thickness at this age. However, the most noticeable difference in the developmental pattern was the shift to more proximal targets after 1 month of age. This is seen when comparing both the average postsynaptic distance of all targets from the soma (Figure 5A), as well as the proportion of synapses located on the soma and proximal dendrites within 20 μm (25 ± 20% at P14 to P30, vs. 63 ± 26% at 2 and 7 months, p = 0.001). Because the postsynaptic neuron in 3 of the pairs at the older ages was not a pyramidal neuron, but another interneuron, we repeated the analysis, now excluding these interneuron-on-interneuron pairs. This did not significantly change the proportion of synapses located on the soma and proximal dendrites within 20 μm at 2 and 7 months (59 ± 32%) nor the significance of the difference with the younger ages (p = 0.01).

**Figure 5.**
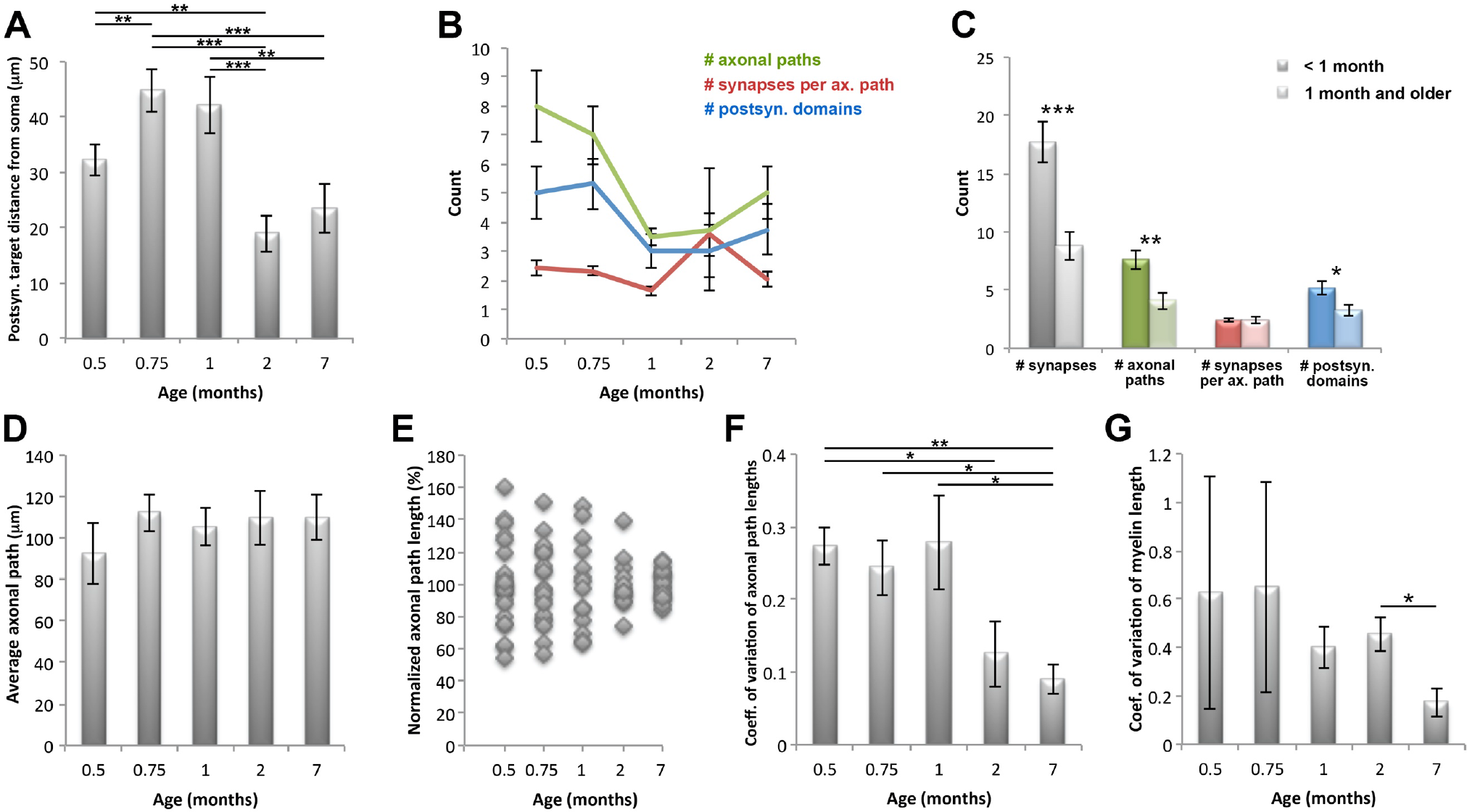
Developmental changes in the location of postsynaptic targets and axonal path lengths in the PV+ interneuron connections. **A.** Changes in postsynaptic target distance from soma. **B.** Changes in the number of axonal paths in a connection, the number of synapses per axonal path, and the number of targeted postsynaptic domains**. C.** Summary of postnatal changes in the number of synapses, axonal paths and targeted postsynaptic domains**. D.** Changes in the average axonal path per connection. **E.** Axonal path lengths normalized to the average per connection. While the variations in the average axonal path length between pairs doesn’t change much, as seen from the error bars in D, the length of individual axonal paths within a connection becomes much more uniform with age. **F.** Changes in the coefficient of variation of axonal path lengths. **G.** Changes in the coefficient of variation of the length of myelinated internodes of axonal paths. For all plots statistical significance is **\***P ≤ 0.05, **\*\***P ≤ 0.01. **\*\*\***P ≤ 0.001.

Why does the change in the postsynaptic distribution of synapses lag behind synapse pruning? Our results suggest that initially, the distance from the postsynaptic soma is not a factor determining which synapses are removed, because at 1 month of age, when synapse number is reduced by two thirds, the distribution of postsynaptic targets remains the same. The subsequent decrease in the postsynaptic target distance from the soma is likely a result of continuing synapse turnover with preferential removal of more distal synapses on dendrites and addition of new somatic / peri-somatic synapses.

### Synapse pruning likely occurs by removal of distal axonal branches

PV+ interneurons send multiple axonal paths to form synapses with the postsynaptic neuron. These axonal paths diverge early on, most commonly starting at the first axonal branch point, and continue to split with increasing axonal branch orders. At P14 there were on average 8 ± 2 separate axonal paths (separated by 2 or more branch points), and at P22, 7 ± 2 axonal paths (Figure 5B). Each of these axonal paths made on average 2.4 ± 0.4 synapses, however, occasionally as many as 9 synapses per axonal path were observed (e.g pair MK190307). This raises the question: does synaptic pruning of PV+ interneurons involve removing of entire axonal branches similarly to what happens with thalamocortical axons and axons of Cajal-Retzius interneurons in developing neocortex (Portera-Cailliau et al., 2005) and at the neuromuscular junction (Tapia et al., 2012), or are only individual synapses eliminated? The number of first and second order axonal branches in a connection remained stable during development (1^st^ branch: 1.9 ± 0.4 at <1 month vs. 1.8 ± 0.6 at 1 month and older; 2^nd^ branch: 3.0 ± 0.6 at <1 month vs. 2.5 ± 0.9 at 1 month and older). However, the number of separate axonal paths (separated by 2 or more branch points) decreased in half from 7.6 ± 2.1 at <1 month to 4.1 ± 2.5 at 1 month and older (p = 0.006, Figure 5B,C). At the same time, the number of synapses per axonal path did not change, strongly suggesting that synaptic pruning predominantly removes entire distal axonal branches and not individual synapses. This is consistent with our observation that the density of synaptic varicosities along axons does not change after P22 (Figure 4C). On the postsynaptic side, this results in fewer postsynaptic domains (cell body, primary dendrite with it branches, or axon initial segment) receiving synaptic input (5.1 ± 1.6 at <1 month vs. 3.2 ± 1.6 at 1 month and older, p = 0.02).

### Prolonged period of connection refinement results in more uniform axonal path lengths and myelination in the mature animal

Does the axon arbor continue to change after the initial period of synapse pruning (P22 – 1month)? The average axonal path length (the average of the connection averages) remained constant throughout development, at 106 ± 21 μm (Figure 5D). However, there was a remarkable change in the variability of the length of the axonal paths in a connection. At the youngest age, P14, the lengths of the different paths in a connection varied by as much as 60% from the average, and similar high variability persisted at P22 and at 1 month with path lengths varying as much as 50% from the average. After 1 month, the lengths of the different paths comprising the connection become much more uniform, and at 7 months axonal path lengths deviated by less than 15% from the average length for the connection (Figure 5E). This is reflected in the significant decrease in the coefficient of variation of axonal path lengths at 2 and especially 7 months (Figure 5F). At the same time, the extent of myelination of the different axonal paths in a connection became much more consistent, and there was a significant reduction in the coefficient of variation of the length of myelinated internodes along axonal paths between 2 and 7 postnatal months (Figure 5G).

### Autapse numbers remain stable between P14 and 7 months

PV+ interneurons also form autapses, i.e. synapses onto themselves, shown to have important functional roles, such as regulating spike-timing precision (Bacci and Huguenard, 2006) and tuning to gamma oscillations (Deleuze et al., 2019). Examples of autapses are shown in Figure 6A,B and E. Even though both the axon and its postsynaptic target in this case are filled with the same dye, the fluorescence in the axon is weaker and thus can be detected as a separate entity when analyzing the images. Autapses are also known to have large presynaptic varicosities (Tamás et al., 1997b), further aiding in their identification. At P14 and P22 autapses were observed in all the neuronal pairs (n=7) and their number ranged from 4 to 12 per interneuron, with an average of 7 ± 3. The number of autapses per interneuron was significantly less than the number of synapses the interneuron made onto the postsynaptic neuron (Figure 6C). Compared to synapses at these ages, they were distributed more proximally to the soma (27 ± 23 μm vs. 37 ± 25 μm, p = 0.03). At later ages, the number of autapses did not change significantly and, in contrast to synapses, no evidence of pruning was observed. Autapses were distributed further away from the soma at 1 and 7 months, but not at 2 months (Figure 6D). No correlation was seen between the number of autapses and the number of synapses per connection, neither between the distribution of autapses and synapses relative to the soma (Figure 6F,G).

**Figure 6.**
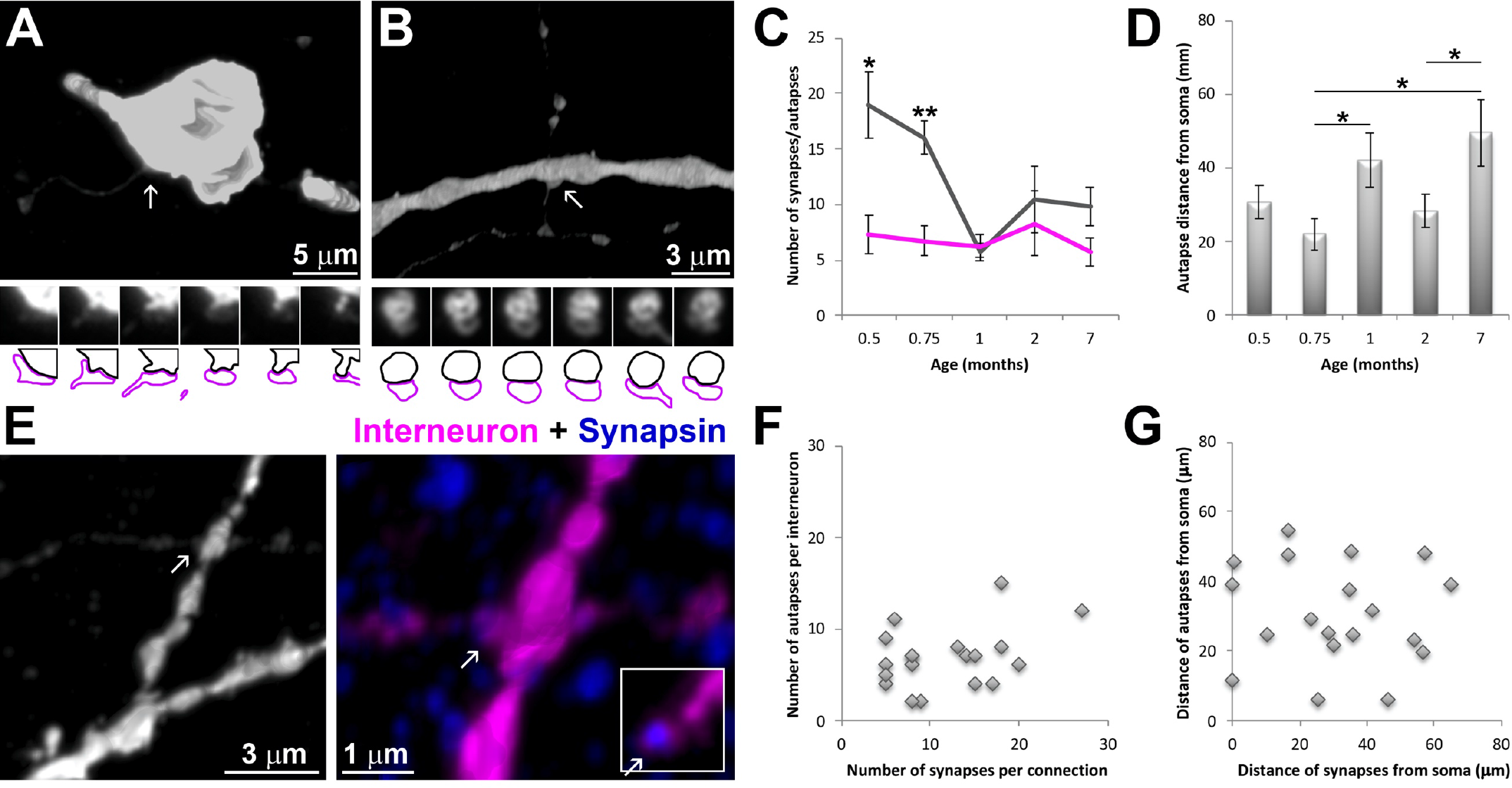
PV+ interneuron autapses. **A.** Example of an autapse on the interneuron soma (2 month old pair MK190115-1). Volume reconstructions of the interneuron (streptavidin Alexa594) on top and serial sections through the autapse on the bottom. The fluorescence intensity of the axon is much weaker compared to that of the soma or dendrites. In the color map of the autapses, the presynaptic bouton is outlined in magenta and the postsynaptic target in black. **B.** Example of an autapse on a proximal dendrite (1 month old pair MK181212). **C.** Developmental changes in the number of autapses per PV+ interneuron compared to changes in the number of synapses. **D.** Developmental changes in the autapse target distribution. **E.** Example of an autapse (white arrow) at P14 (pair MK190405-1). Volume reconstruction of the interneuron only (left, Streptavidin Alexa594) and zoomed in view of the synapse with the interneuron in magenta and synapsin immunolabel in blue (right). The insert to the right is a single section through the autapse showing the colocalization of synapsin (blue) and the presynaptic bouton fluorescence (magenta). **F.** The number of autapses does not correlate with the number of synapses in the mapped connections made by the PV+ interneurons. **G.** There is no correlation between the target distribution of the autapses and the synapses made by the PV+ interneurons.

## Discussion

PV+ basket cells in mammalian neocortex have distinct axonal arbors, characterized by extensive branching, intermittent myelination, and abundant synapses surrounding cell bodies of neighboring pyramidal neurons and within the neuropil. Our results reveal the extensive reorganization of the axon-myelin unit following the formation of the initial immature connectivity of PV+ interneurons (Figure 7). At the end of the second postnatal week, PV+ basket cells have highly branched axons that are rarely myelinated and form abundant synaptic contacts with pyramidal neurons. During the third postnatal week the proximal axons of PV+ interneurons become rapidly myelinated, and in the fourth postnatal week massive synapse pruning occurs. This pruning does not affect the autapses that PV+ interneurons form on themselves. More subtle reorganizations continue throughout the examined period, until postnatal month 7. They include a shift of the postsynaptic targets of PV+ interneurons to more proximal locations and a significant decrease of the variability of the length of axonal paths in each connection, as well as continued myelination of the translaminar axonal branch that extends further away from the interneuron soma.

**Figure 7.**
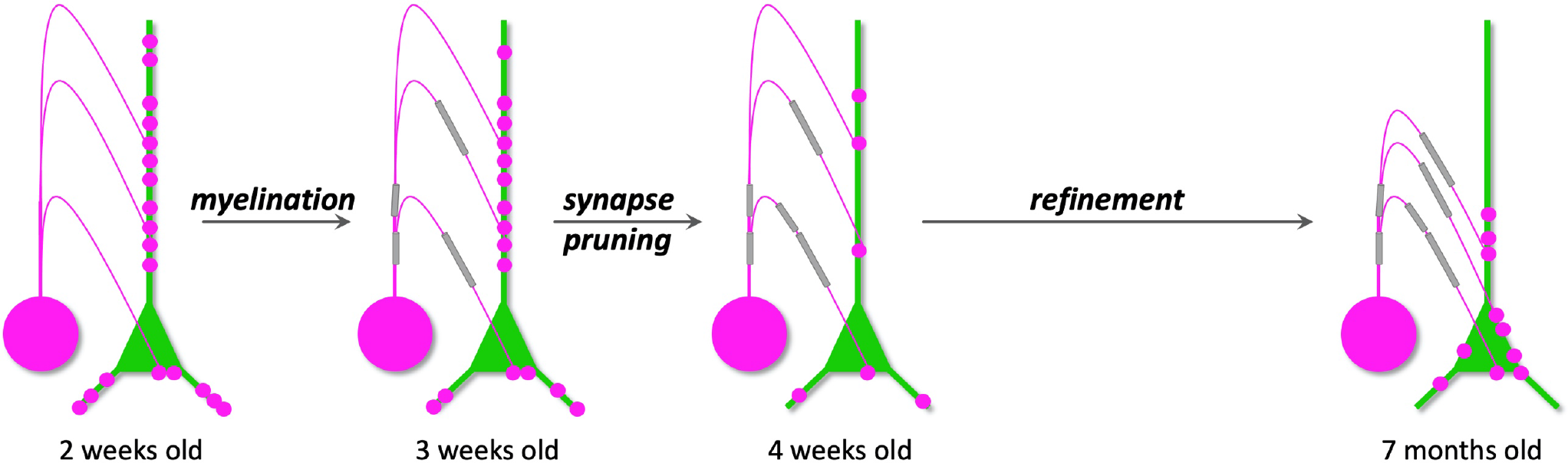
Summary of the main developmental changes in PV+ interneuron individual connections. Axonal myelination develops between the 2^nd^ and 3^rd^ postnatal week, followed by massive synapse pruning resulting in about three-fold reduction of synapses in the PV+ interneuron connections by the 4th postnatal week. Connection refinement continues beyond this, with the postsynaptic targets of PV interneurons gradually shifting to more proximal locations, and the length of axonal paths and their myelin becoming conspicuously uniform per connection by postnatal month 7. PV+ interneuron is in magenta and postsynaptic neuron in green. Myelin is in grey.

### Developmental changes in myelination of PV+ interneurons

Myelination of the axon is a characteristic feature of PV+ basket cells in neocortex (DeFelipe et al., 1986; Somogyi and Soltesz, 1986; Micheva et al., 2016; Stedehouder et al., 2017), despite the fact that the great majority of their connections are local (Packer et al., 2011). Deficits in myelination can affect multiple aspects of PV+ interneuron development, such as their axonal arborization, synaptic connectivity, and firing rate (Benamer et al., 2020). PV+ interneuron myelin likely has multiple roles, including energy conservation (Hartline and Colman, 2007), and nutrient support (Fünfschilling et al., 2012; Lee et al., 2012) of these fast-spiking neurons known for their high energy demands (Kann et al., 2014), as well as increase in conduction velocity along the axon (Micheva et al., 2021). Our present results show that during development, myelination of the proximal PV+ interneuron axon occurs rapidly during the 3^rd^ postnatal week. This is also the time when the spiking frequency of PV+ interneurons greatly increases (Okaty et al., 2009; Miyamae et al., 2017), resulting presumably in increased energy demands, and basket cells differentiate into ‘fast signaling devices’ (Doischer et al., 2008) needing faster conduction of axon potentials along the axon. By the end of the first postnatal month the extent of myelination of the proximal axonal arbor (up to 150 μm from soma), as well as the myelination of the axonal paths connecting the interneurons with nearby cells, reach adult levels. Meanwhile, the translaminar axonal branch present in many of the interneurons that we analyzed, continues to increase its myelination, reaching 92% of its length beyond the axon initial segment by 7 months. This suggests that translaminar PV+ interneuron connectivity, as opposed to the intralaminar connectivity explored in this study, may follow a different, delayed developmental trajectory. Interestingly, the extent of myelination of individual PV+ interneurons varies greatly at all ages and we recently showed that it correlates with the conduction velocity along the axon (Micheva et al., 2021). It remains to be seen whether it also correlates with the activity level of interneurons, and, correspondingly, their energy demands.

### PV+ interneuron synapse pruning

Following the initial rapid myelination of PV+ interneuron axons in the 3^rd^ postnatal week, by the end of the first postnatal month the number of synapses per connection declines to one third of the P14 numbers. The observed large drop in the number of synapses per connection was rather surprising, because even though pruning of PV+ interneuron synapses is known to occur at this developmental stage, it is counterbalanced by new synapse formation (Chattopadhyaya et al., 2004). Thus, during the second half of the first postnatal month, there is no significant change in the number of PV+ perisomatic synapses in layer 2/3 of rat medial entorhinal cortex (Berggaard et al., 2018) and an increase in perisomatic PV+ synapses in mouse visual cortex organotypic cultures (Chattopadhyaya et al., 2004), as well as an increase in the total number of all inhibitory synapses (Micheva and Beaulieu, 1996; DeFelipe et al., 1997). Our findings suggest that the overall increase in PV+ synapse numbers in cortex during the first postnatal month is most likely due to PV+ basket cells continuing to extend their axons and contacting more and more postsynaptic neurons, thus masking the drop in the number of synapses per individual connection. The decrease in PV+ synapses per connection appears to be due to retraction of distal axonal branches, and not removal of individual synapses. Indeed, as shown in organotypic slices from mouse visual cortex (Wu et al., 2012), the axons of PV+ basket cells at this age are very dynamic. New axonal branches continuously emerge and form synapses, while existing axonal branches retract, and branch addition is slightly more prevalent than branch elimination. Synapse elimination is regulated by activity dependent mechanisms, including GABA neurotransmission (Wu et al., 2012). Interestingly, the number of autapses made by PV+ interneurons does not undergo pruning, suggesting that they follow different developmental rules compared to synapses. While autapse formation and maintenance may be an intrinsic property of PV+ basket cells, synapse formation involves a two-way communication with the postsynaptic neuron. Previous studies have shown that inhibitory synapses undergo pruning, but to a lesser extent than excitatory synapses (DeFelipe et al., 1997). Our results strongly suggest that PV+ basket cells initially form exuberant synaptic connections that are subsequently refined as weaker or inappropriate synapses are removed. The magnitude of this refinement (a three-fold reduction in synapse numbers per connection), however, can be appreciated only when focusing on individual neuron-to-neuron connections as done in the current study, because synapse pruning is masked to a large extent by the formation of new synapses.

### Limitations

There are several caveats to our study to take into account when interpreting the results. Because we used Wisteria floribunda lectin labeling of perineuronal nets to identify PV+ interneurons, and PNN development is activity-regulated (Dityatev et al., 2007), it is possible that this method of identification selected specifically for PV+ interneurons with higher activity levels and/or more advanced maturation at P14 and P22. Consequently, the developmental changes that we report may be underestimating the magnitude of axonal and synaptic reorganization of neocortical PV+ basket cells during the first postnatal month. Additionally, the PV+ interneurons that were analyzed come from different cortical layers (layers 3 through 6) of entorhinal and perirhinal cortex. It is highly likely that this introduces additional variations in their axonal organization and connectivity that may be obscuring some developmental trends. Other important limitations come from our choice of experimental methods. Immunofluorescent array tomography, while providing the resolution to reliably trace PV+ interneuron axons and their myelin sheath, and identify the great majority of synapses, requires chemical fixation and ultrathin sectioning of the tissue, which is incompatible with live imaging. Developmental trajectories are inferred from static images at different time points, which introduces ambiguities. For example, we report that the myelination of the translaminar axon shows a statistically significant increase between 2 and 7 months, but the data is insufficient to draw any conclusions about how PV+ interneuron myelination compares to the overall cortical myelination, which is known to increase throughout much of adulthood (Hill et al., 2018; Hughes et al., 2018).

### Functional implications of the developmental changes of PV+ interneuron connectivity and myelination

The profound changes in axon myelination and synaptic connections of PV+ interneurons during the second half of the first postnatal month coincide with rapid behavioral changes. The PV+ interneurons analyzed in this study were located in entorhinal and perirhinal cortices, higher-order cortical regions involved in spatial navigation (Moser et al., 2008) and object recognition memory (Squire et al., 2007). Mouse pups open their eyes at P14 and begin venturing outside of their nest around P16 (König 2012), and this exploratory activity greatly increases by the end of the third postnatal week, a period that overlaps with the myelination and immediately precedes the massive synapse pruning of individual PV+ connections that we report. Rodent exploration behaviors, however, continue to mature until the third postnatal month (Wills et al., 2014), and consistent with this, we observe more gradual alterations of the PV+ interneuron axonal arbors and synaptic circuitry past 1 and 2 months of age. These late circuit refinements include a shift of the postsynaptic target locations closer to the postsynaptic cell body, thus ensuring stronger inhibitory control of the postsynaptic neuron. It is notable, however, that PV+ basket cell axons have a strong preference for perisomatic postsynaptic targets already at P14 as seen from our results. This preference likely appears much earlier, between P7 and P9, when a subset of inhibitory axons, which presumably includes PV+ basket cell axons, begin exhibiting a preference for soma innervation, as shown by electron microscopic mapping of inhibitory circuits in the developing mouse somatosensory cortex (Gour et al., 2021). During early development, the establishment of this target preference is likely achieved by pruning of more distal synapses, as evidenced by the decrease in the density of such synapses along the soma-preferring axons (Gour et al., 2021). The continued shift to perisomatic synapse locations later in development that we describe here, may be very similar and achieved through ongoing synapse turnover, albeit at lower levels as observed for GABA synapses in general (Marik et al., 2010, Chen et al., 2011), with new synapses formed closer to the postsynaptic cell bodies, and existing more distal synapses eliminated.

Another change in PV+ interneuron connectivity reported here is the rather striking uniformity of the length of the axonal paths within individual PV+ interneuron connection that is achieved later in development, after the first postnatal month. PV+ interneurons typically connect to their postsynaptic target via several different axonal paths (Micheva et al., 2021), which may provide redundancy to ensure reliable neurotransmission by guarding against possible branch failures. Alternatively, this structural arrangement may enable a much more nuanced communication by allowing differential activation of the paths at axonal branch points (Debanne 2004). In either case, because the extent of myelination of the different axonal paths within a connection also tends to become more even with age, these observations reveal that continued microcircuit refinement results in a remarkably precise organization of PV+ interneuron axonal arbors, providing the structural support for their robust inhibitory effect and fine temporal precision. Taken together, these refinements toward uniform path length and myelination suggest both redundancy in the connection leading to an increase in the ‘safety factor’ of PV+ interneuron to target transmission, but also uniform latency of the synaptic transmission at all of a particular afferent cell’s branches.

## Conclusion

In summary, here we have characterized the development of PV+ interneuron connectivity by focusing on the minimal element of the neuronal circuitry, one neuron synapsing onto one other neuron. Essential for this analysis is the comprehensive investigation of the axon-myelin unit (Simons et al., 2014, Suminaite et al., 2019) and the inclusion of the post-synapse as part of this unit. Our study reveals the elaborate organization and fine precision of the mature PV+ interneuron connectivity and how it evolves during development. Future studies are needed to unravel the complex interactions between the presynaptic and postsynaptic neuron, and the myelinating oligodendrocytes, that underlie these changes.

## Extended Data

**Figure 3-1.**
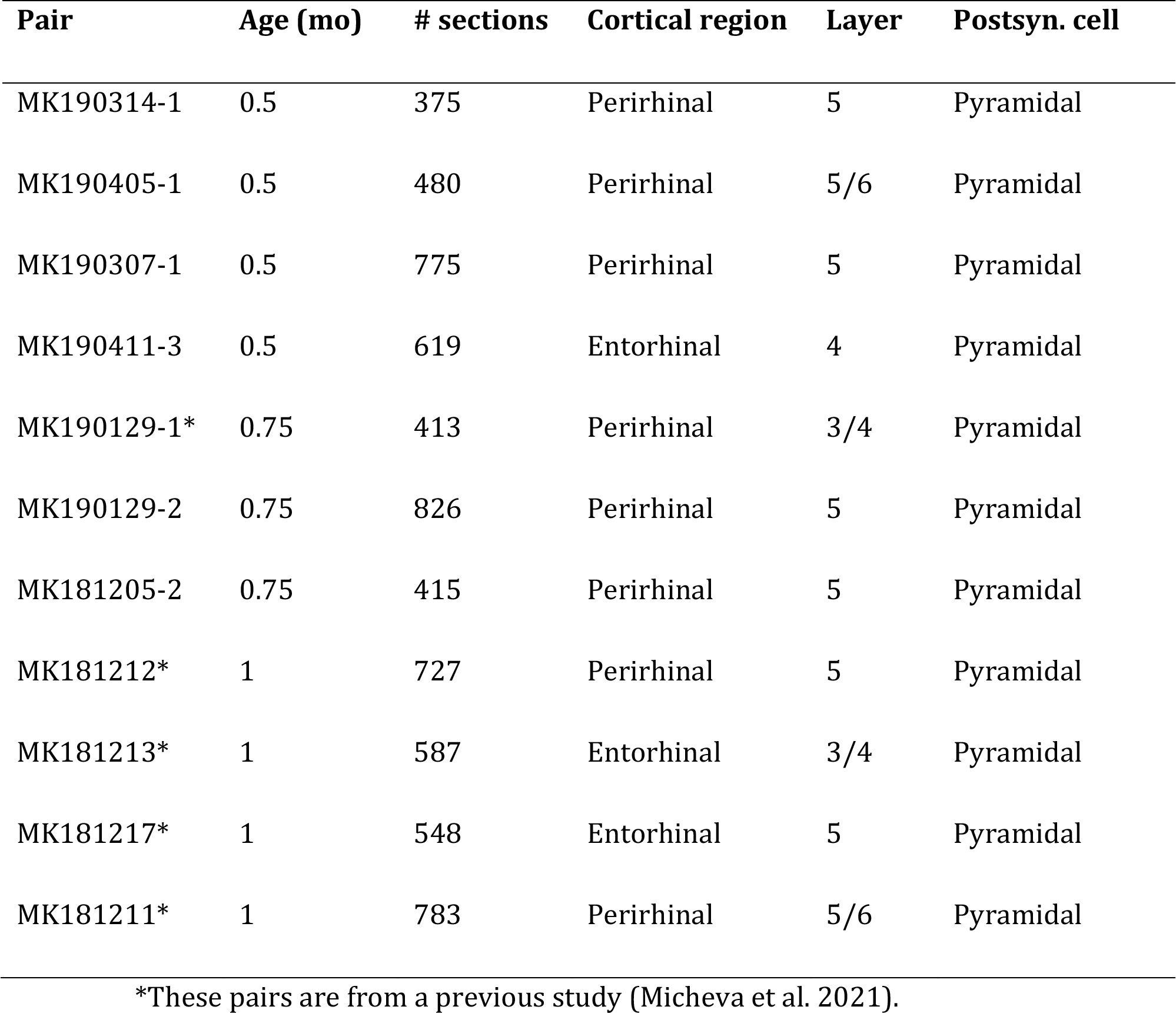
Neuronal pairs analyzed by AT.

**Figure 3-2.**
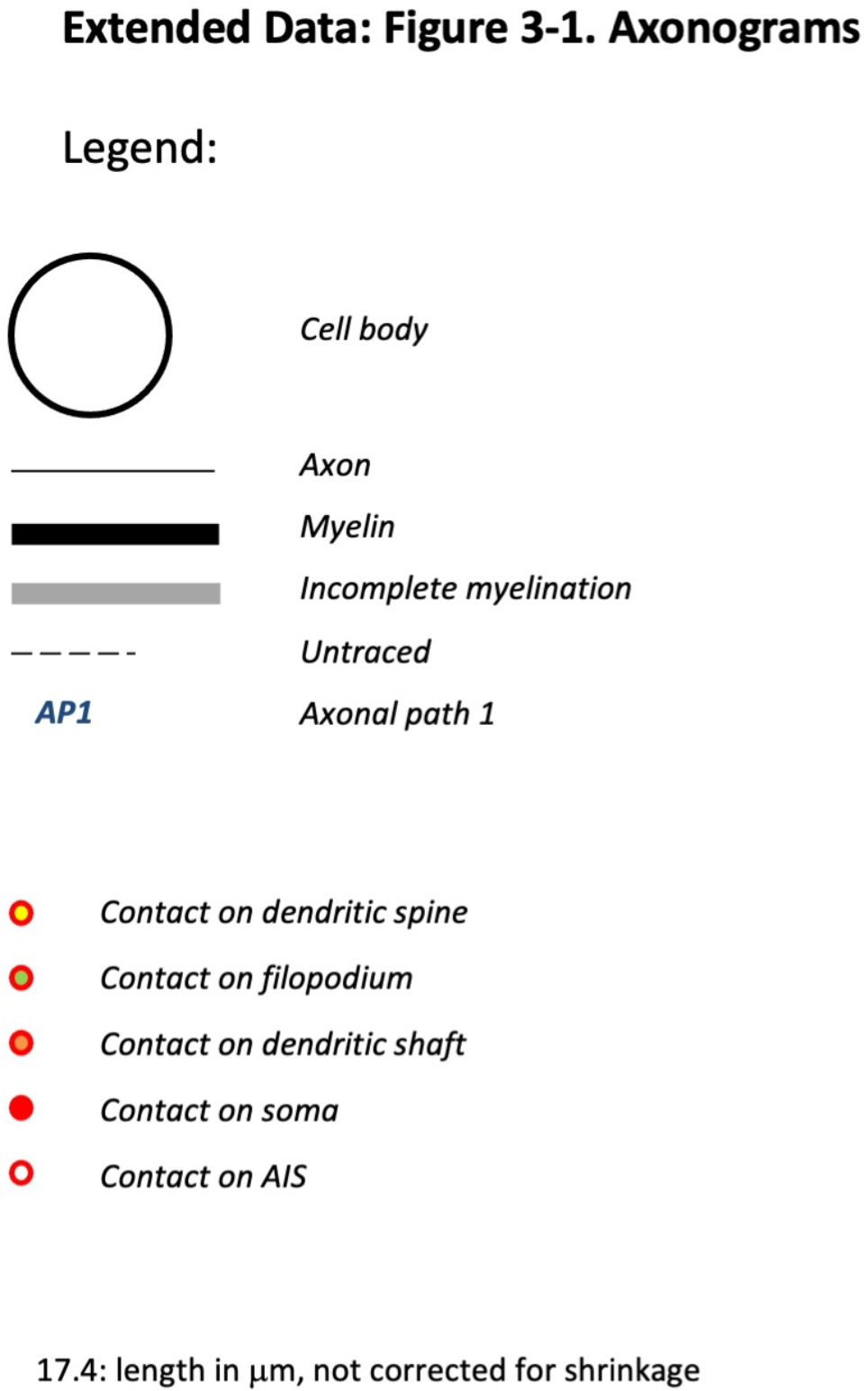
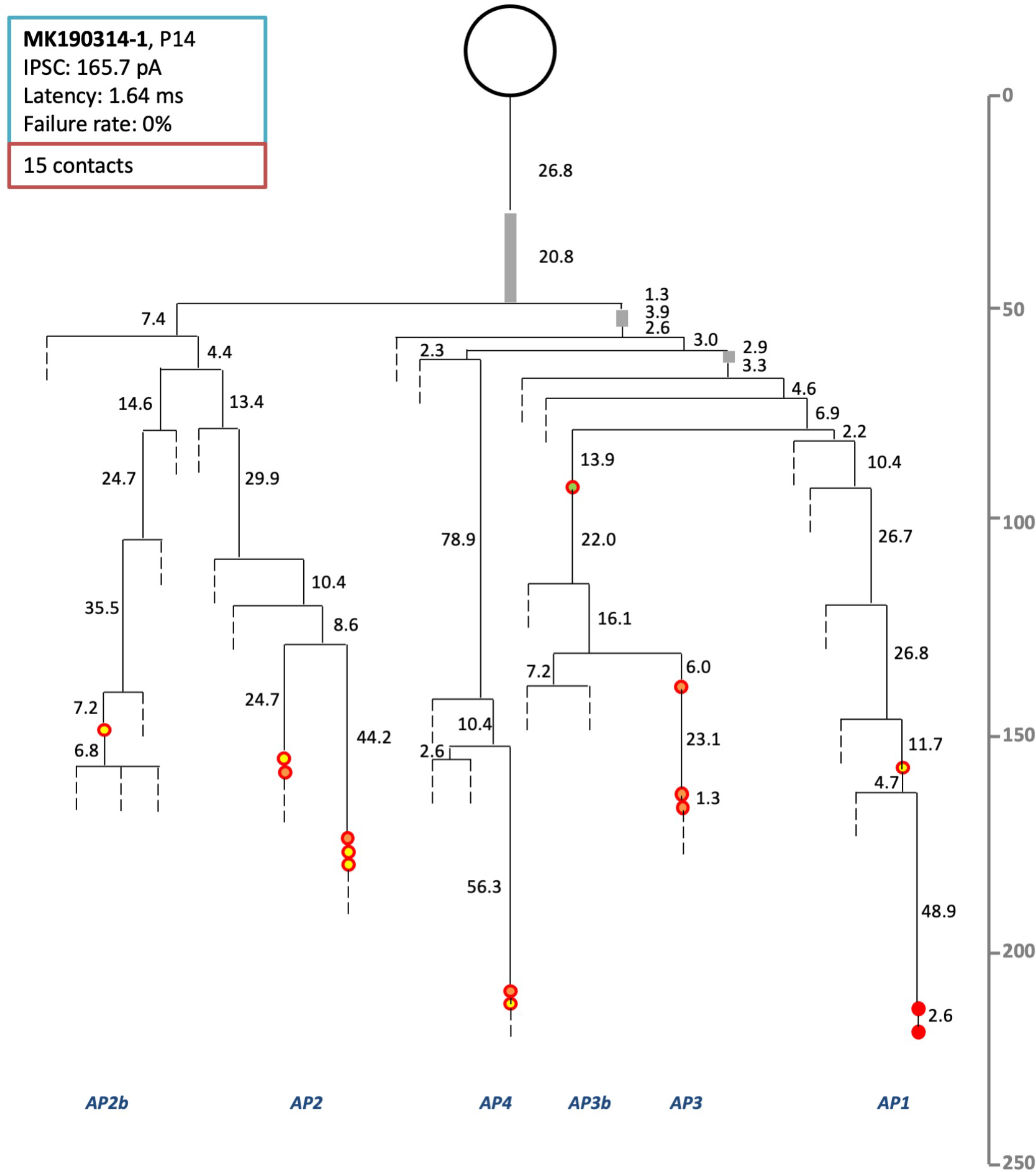
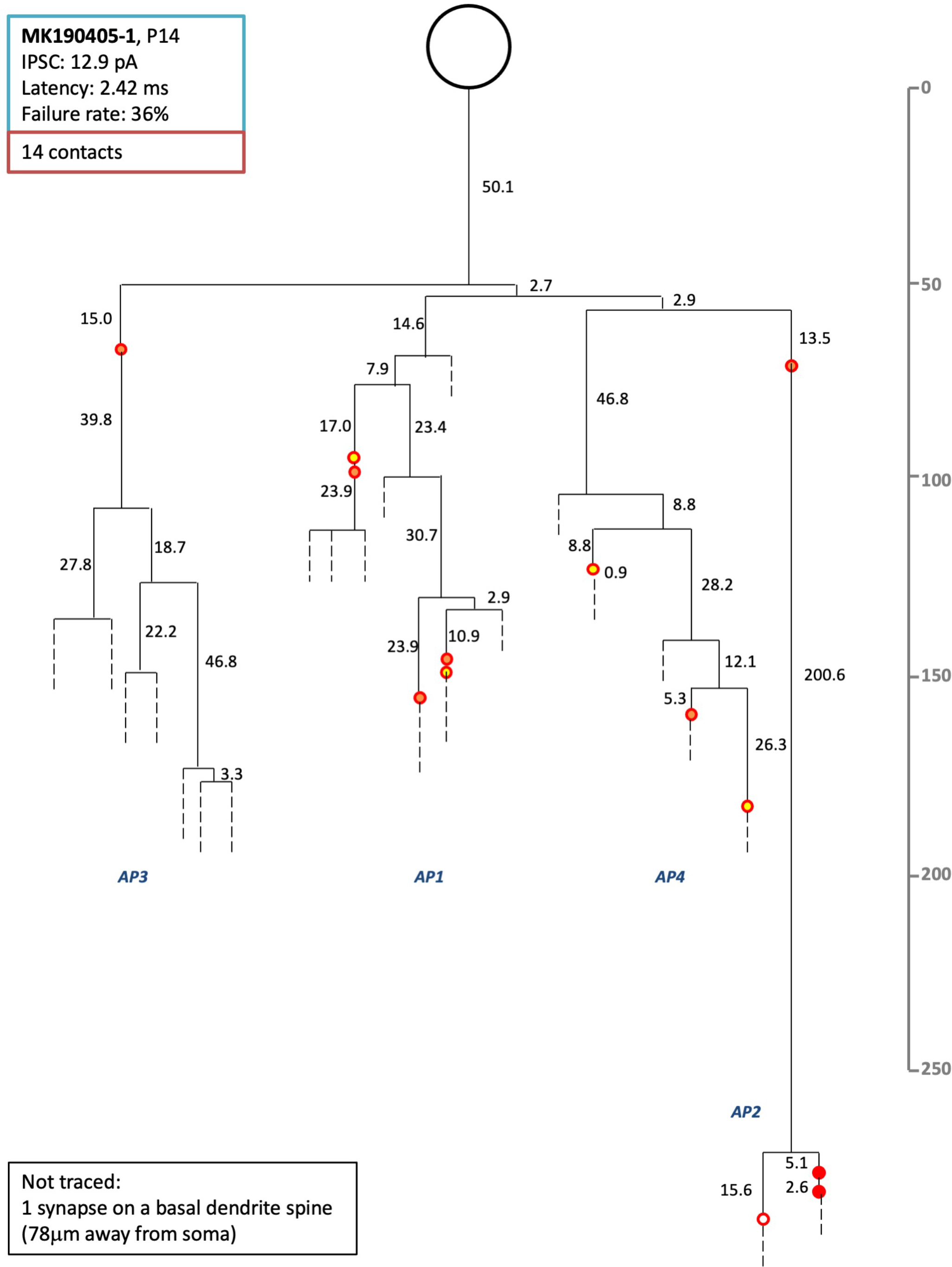
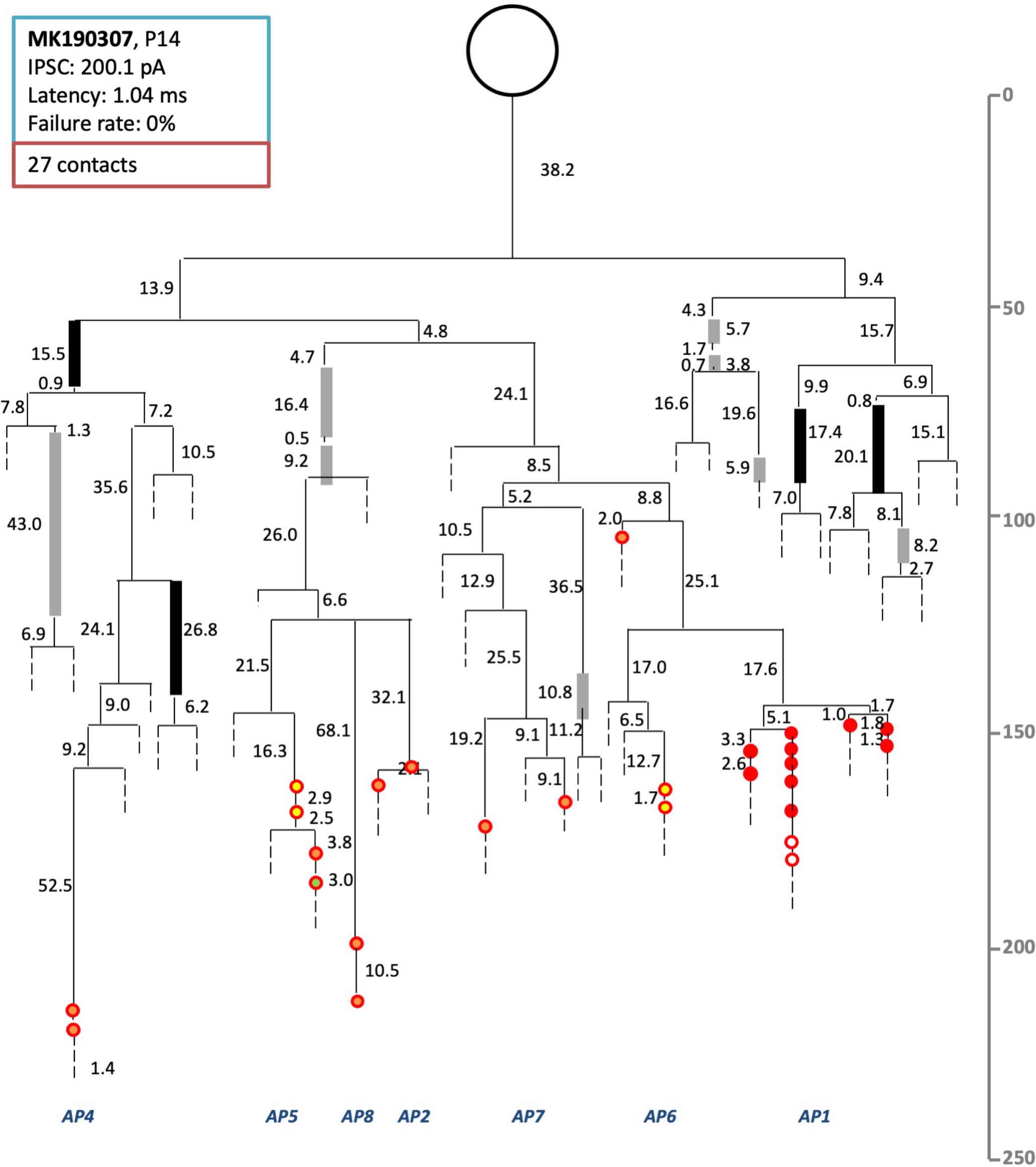
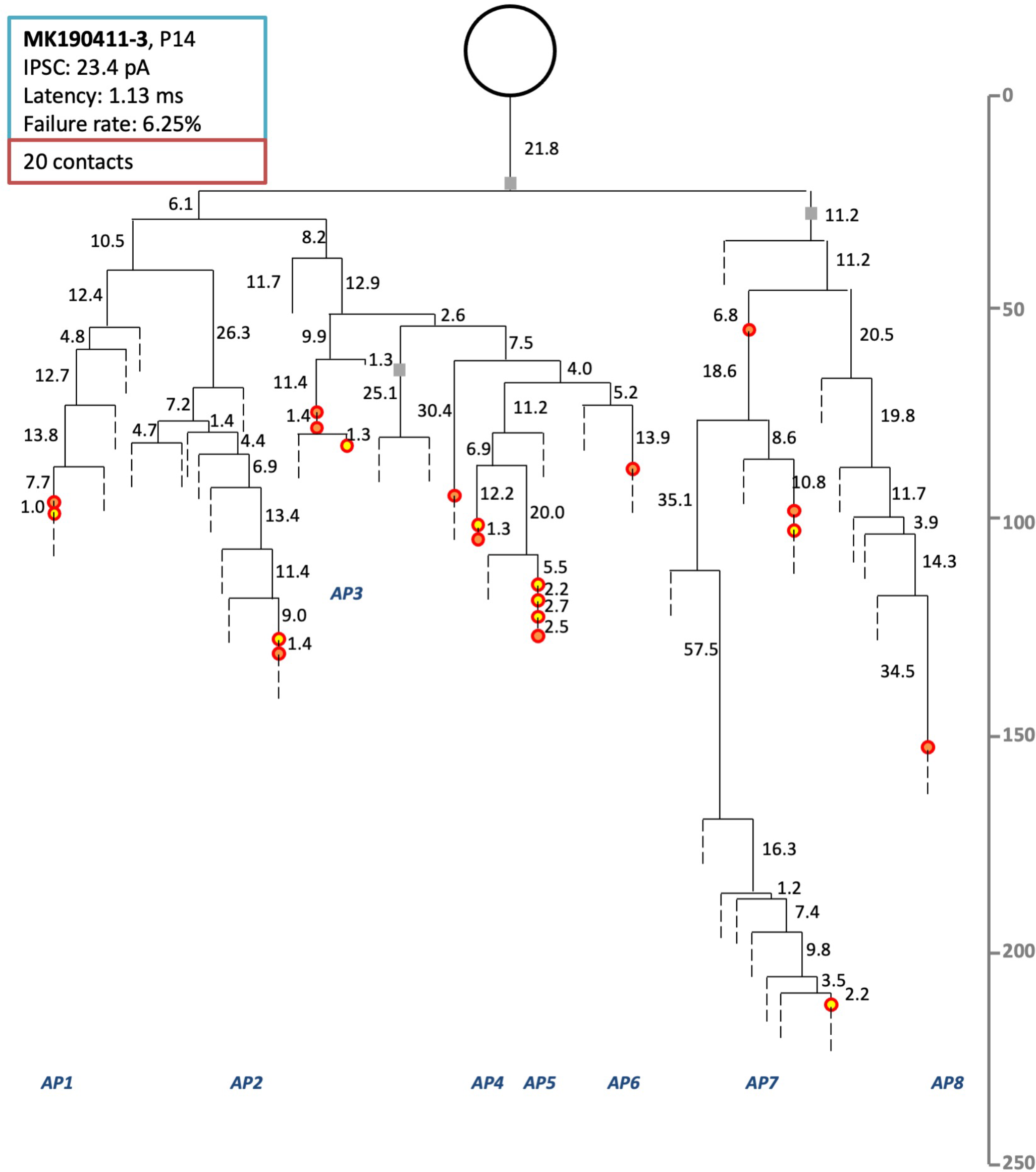
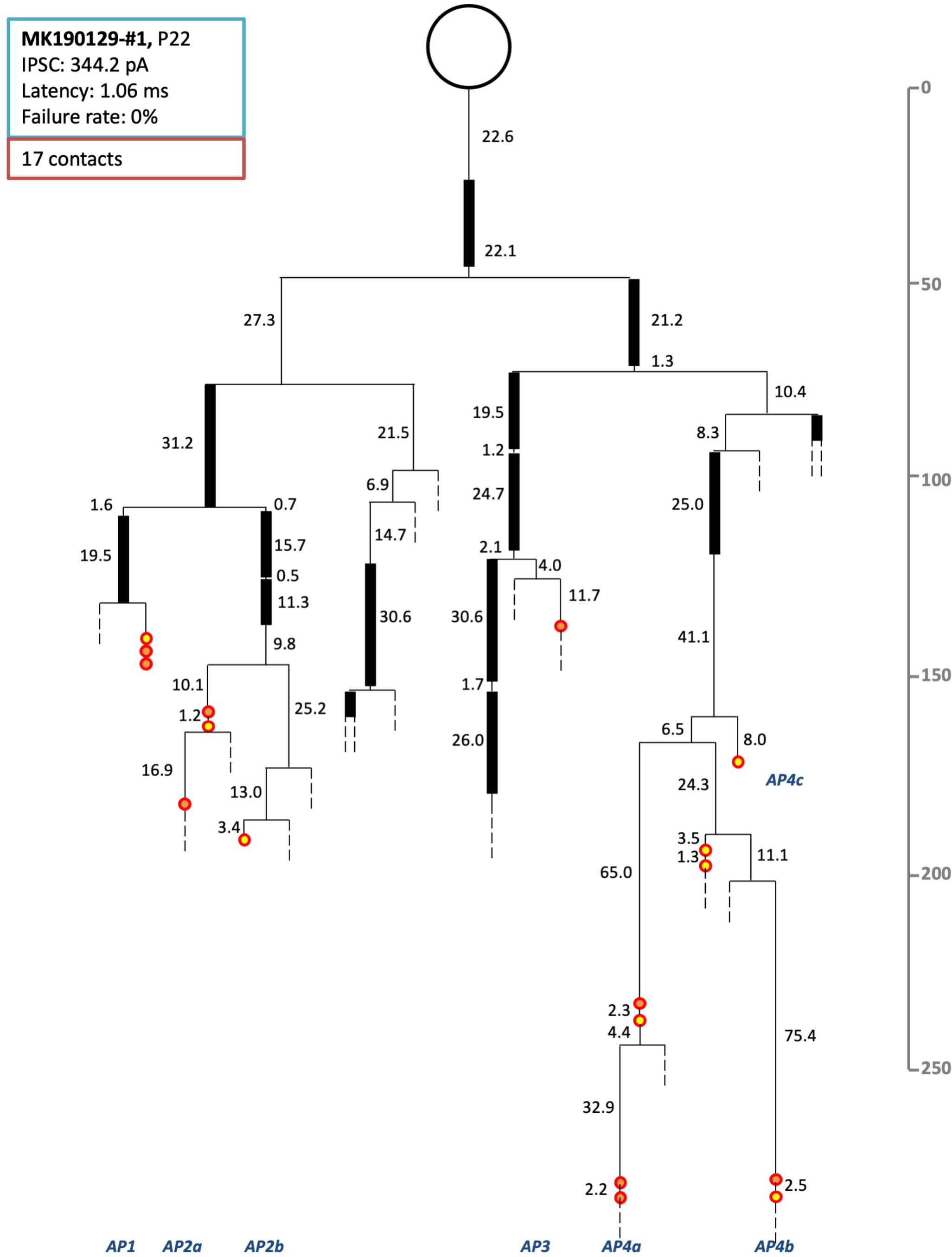
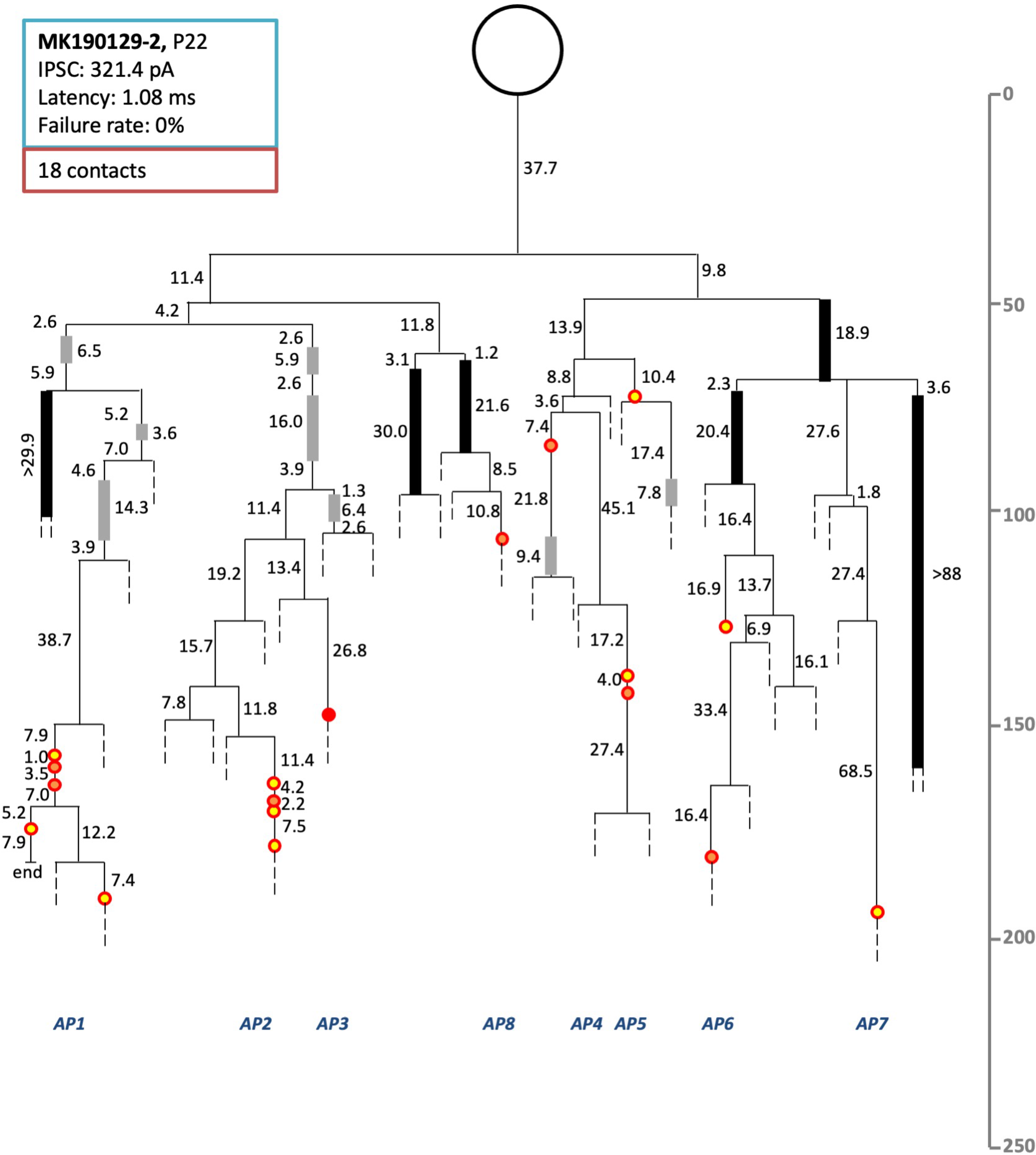
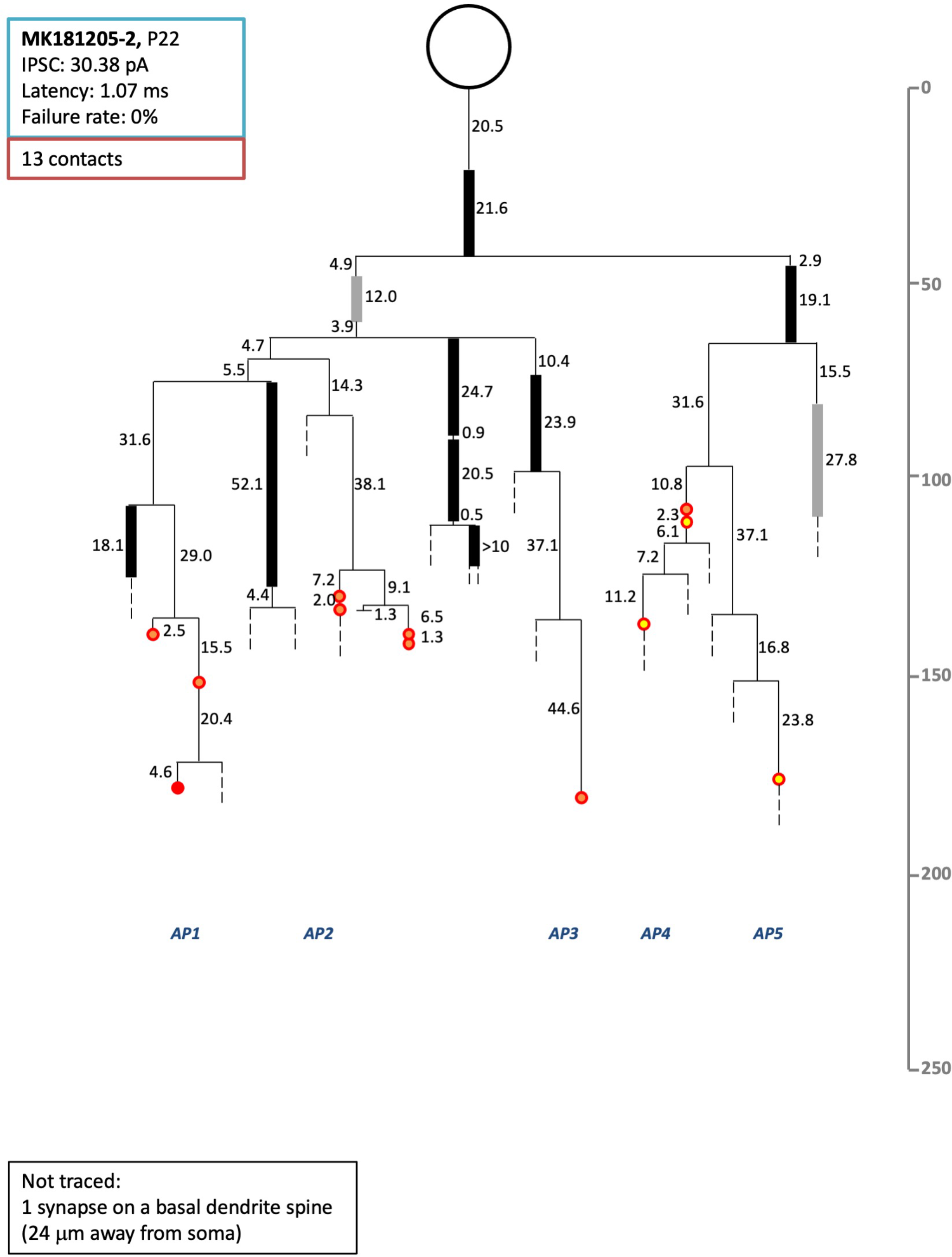
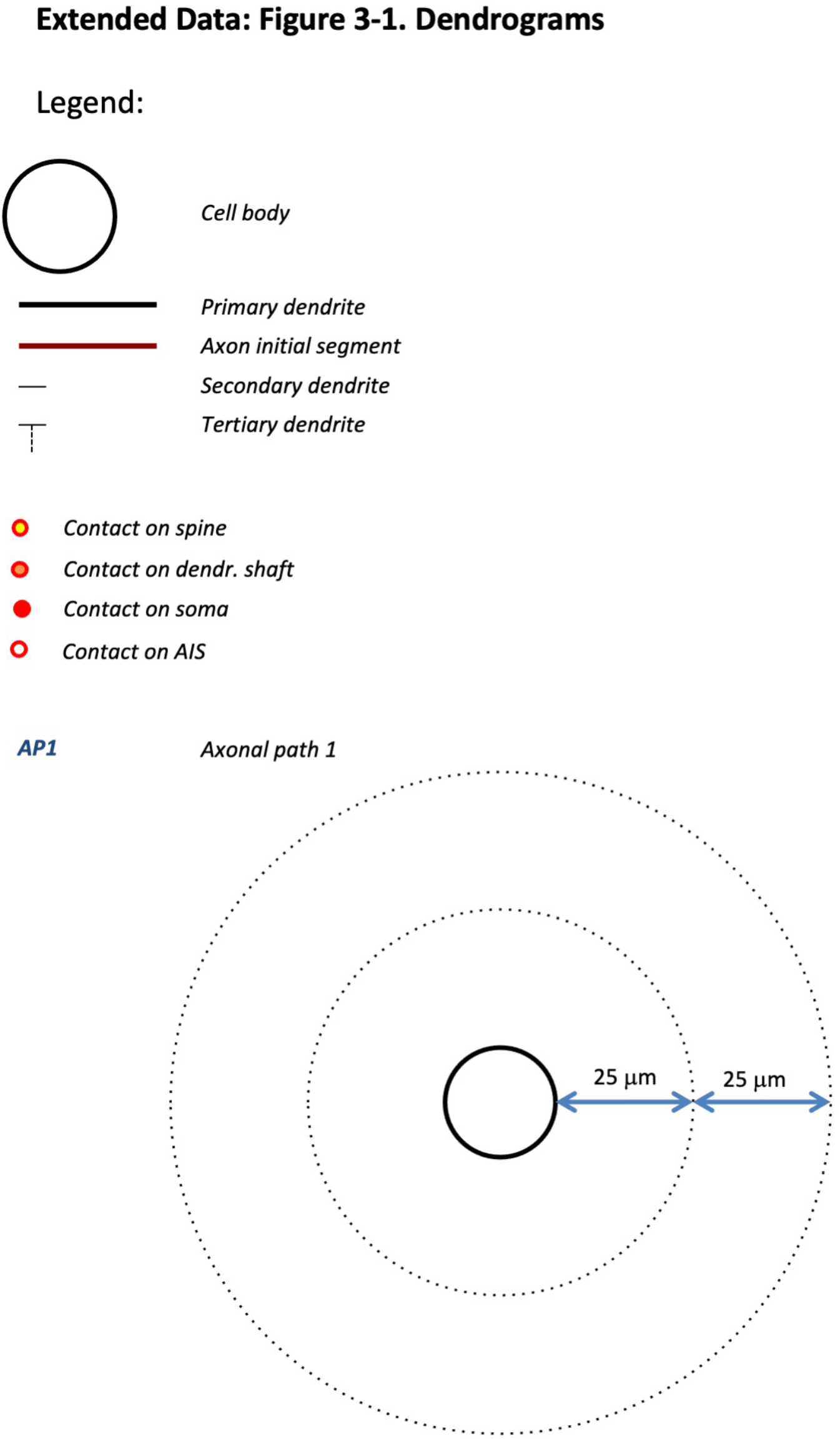
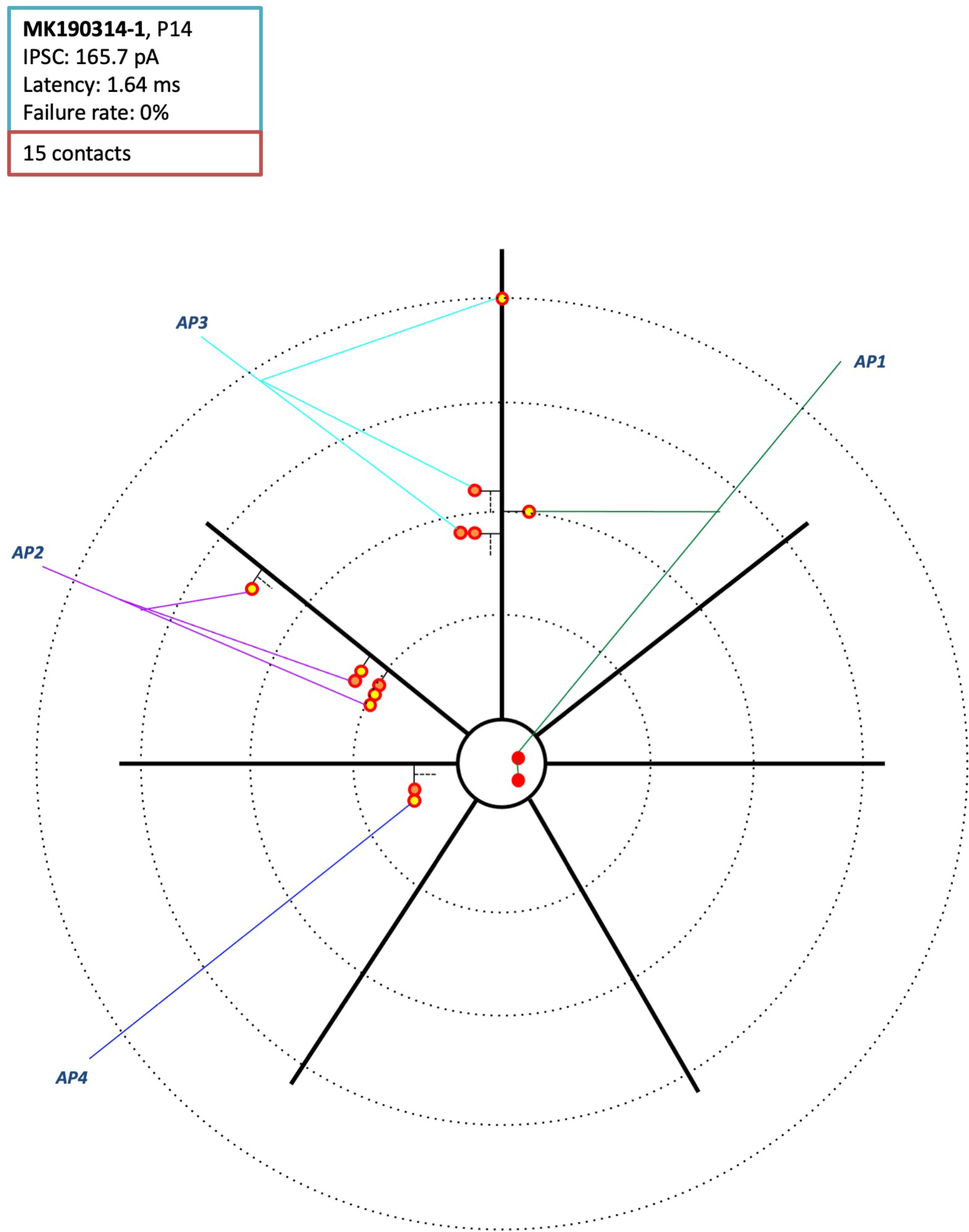
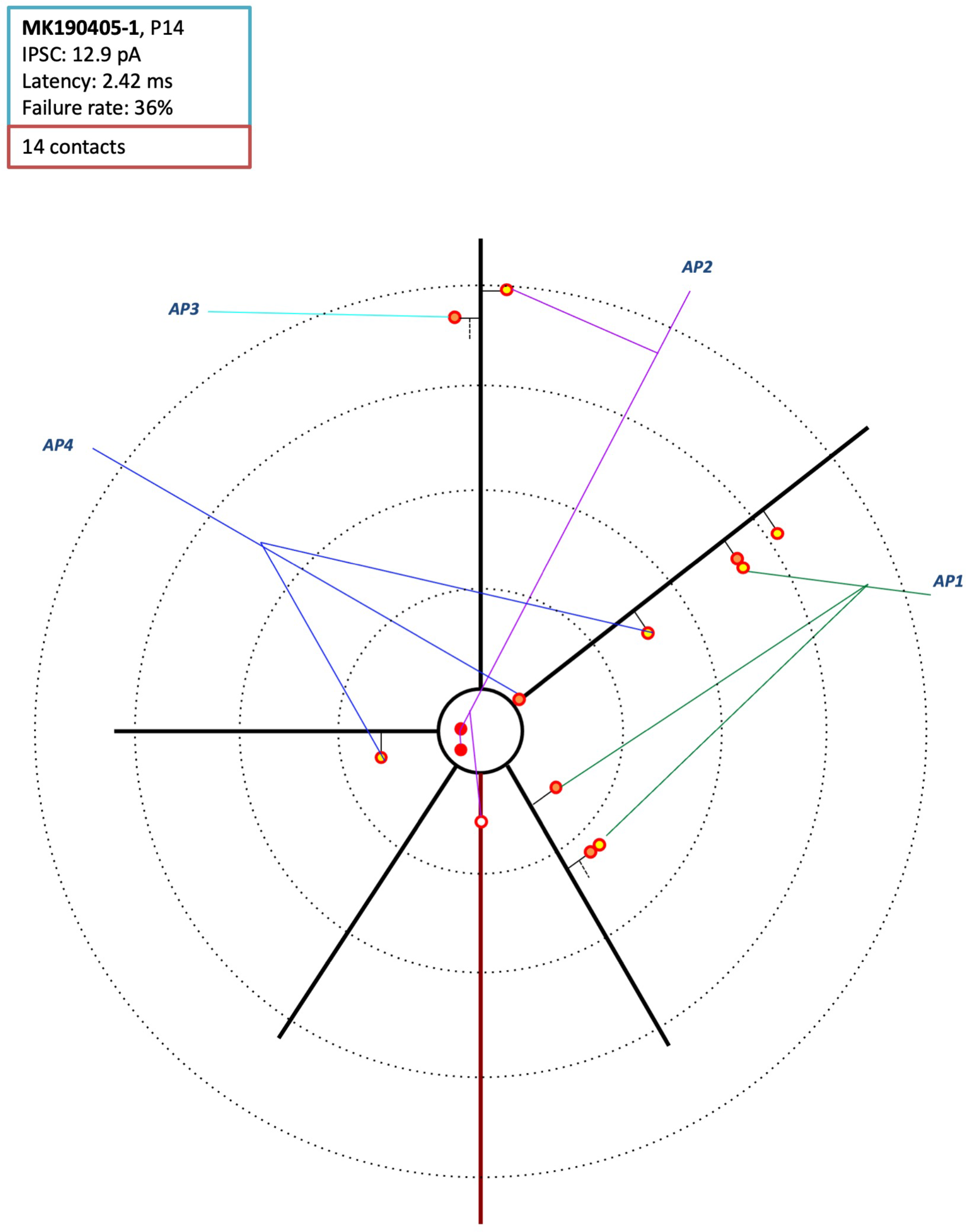
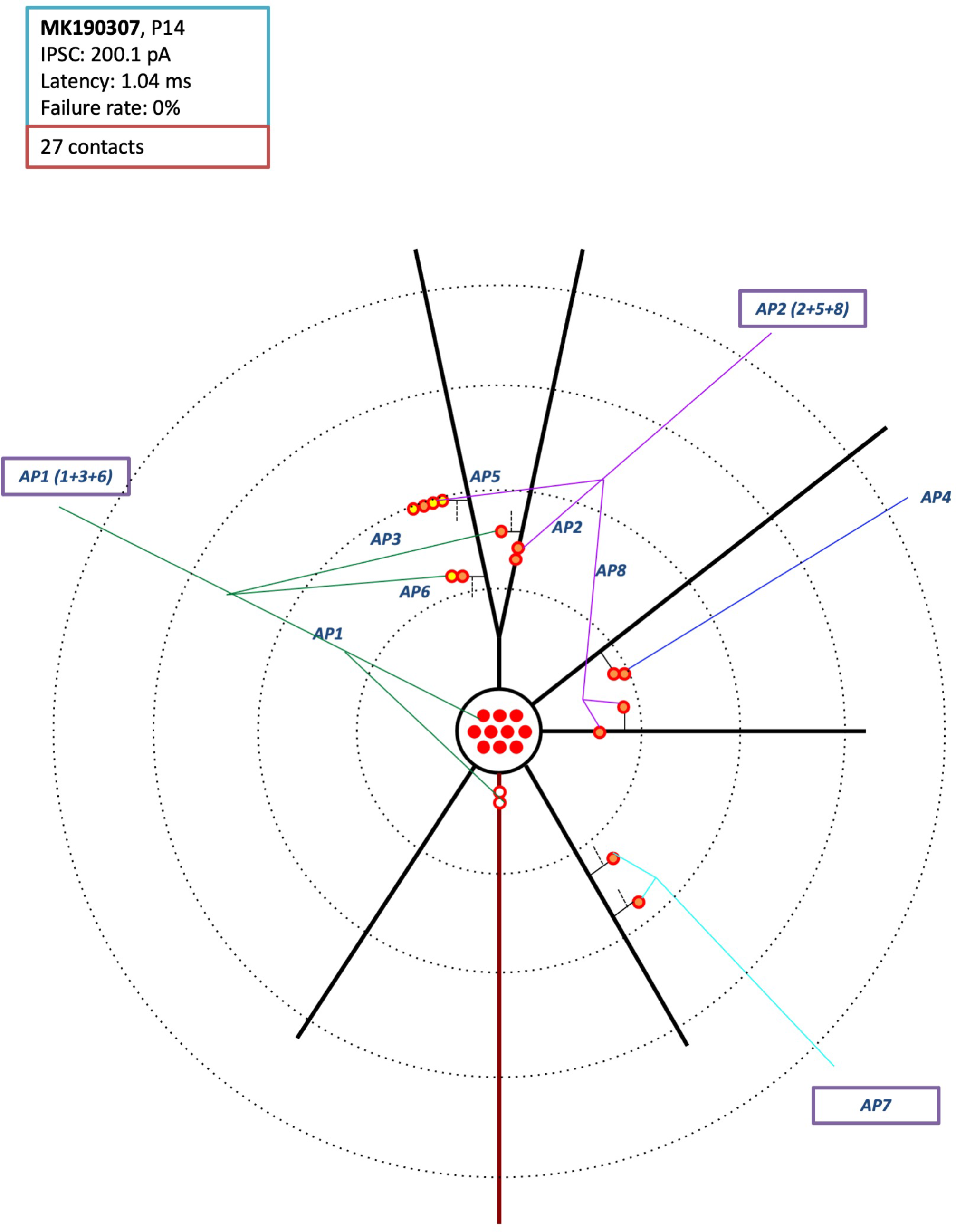
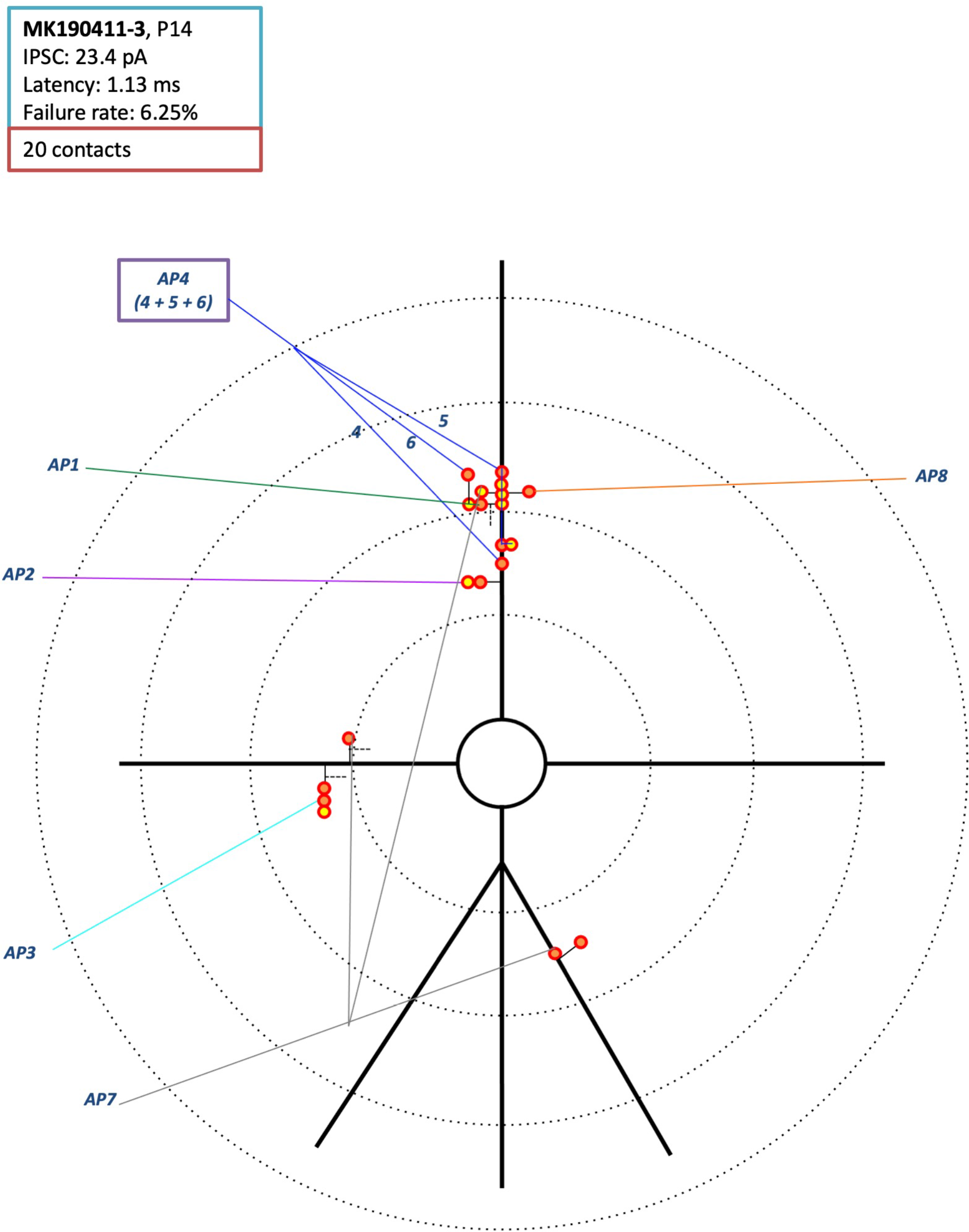
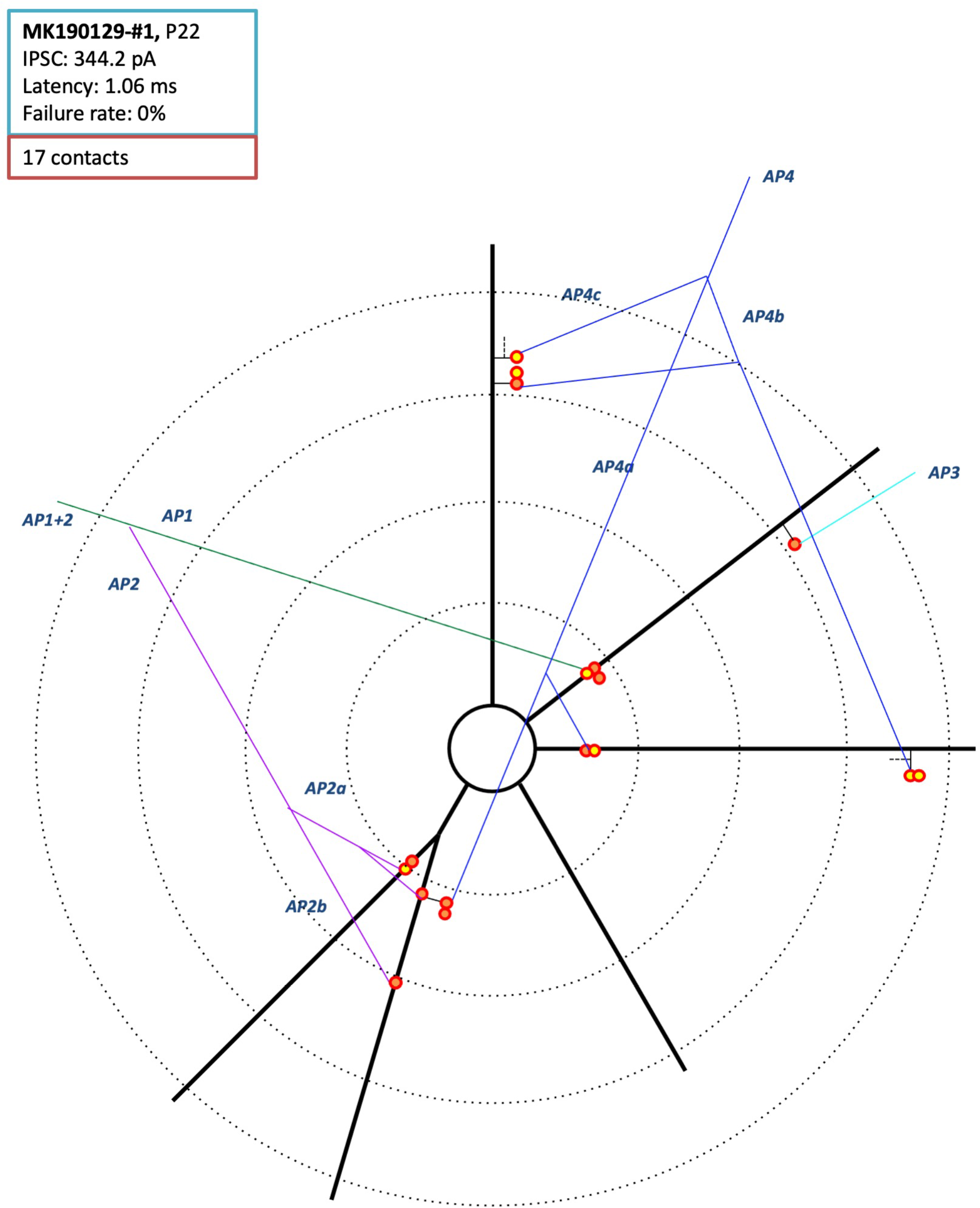
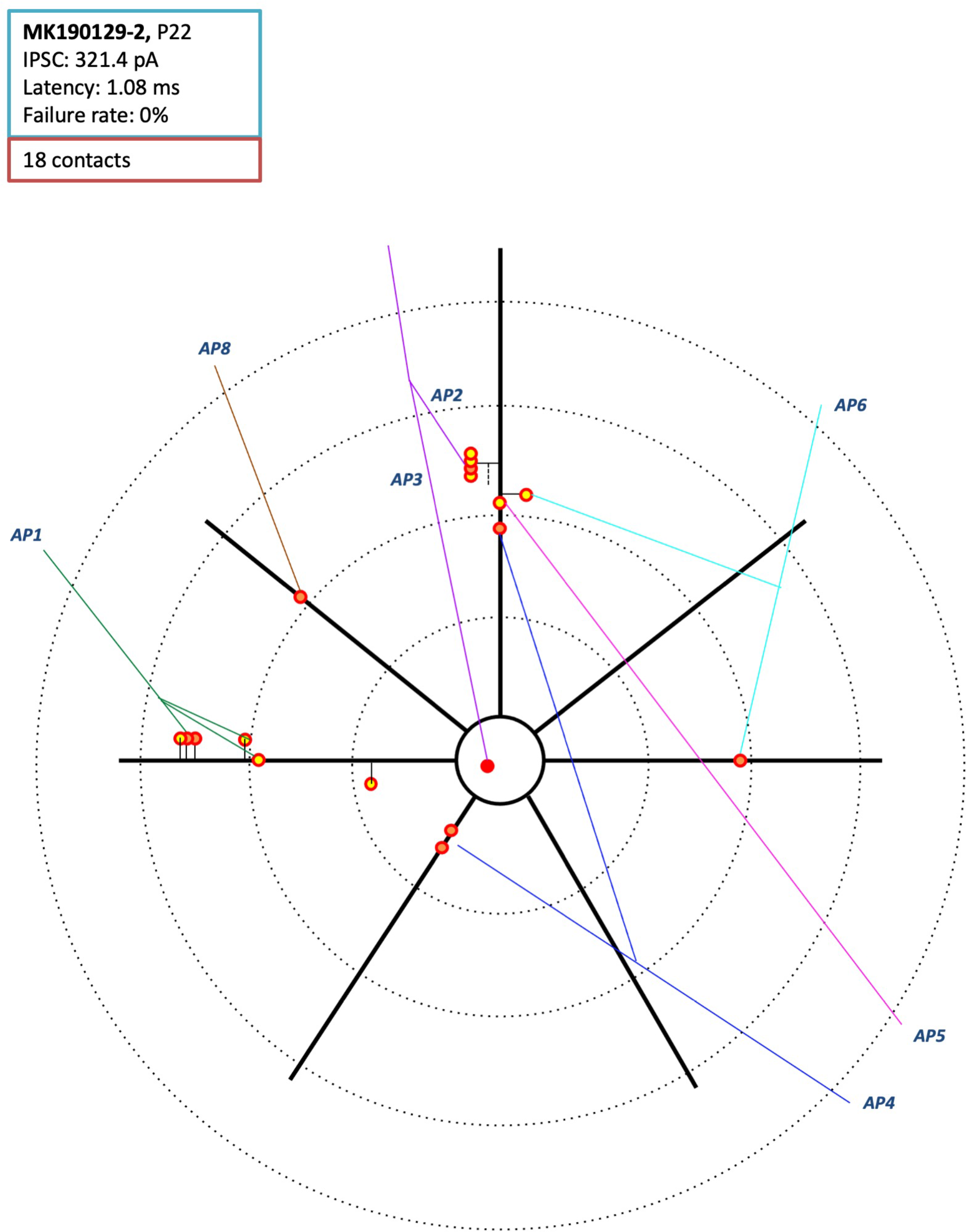
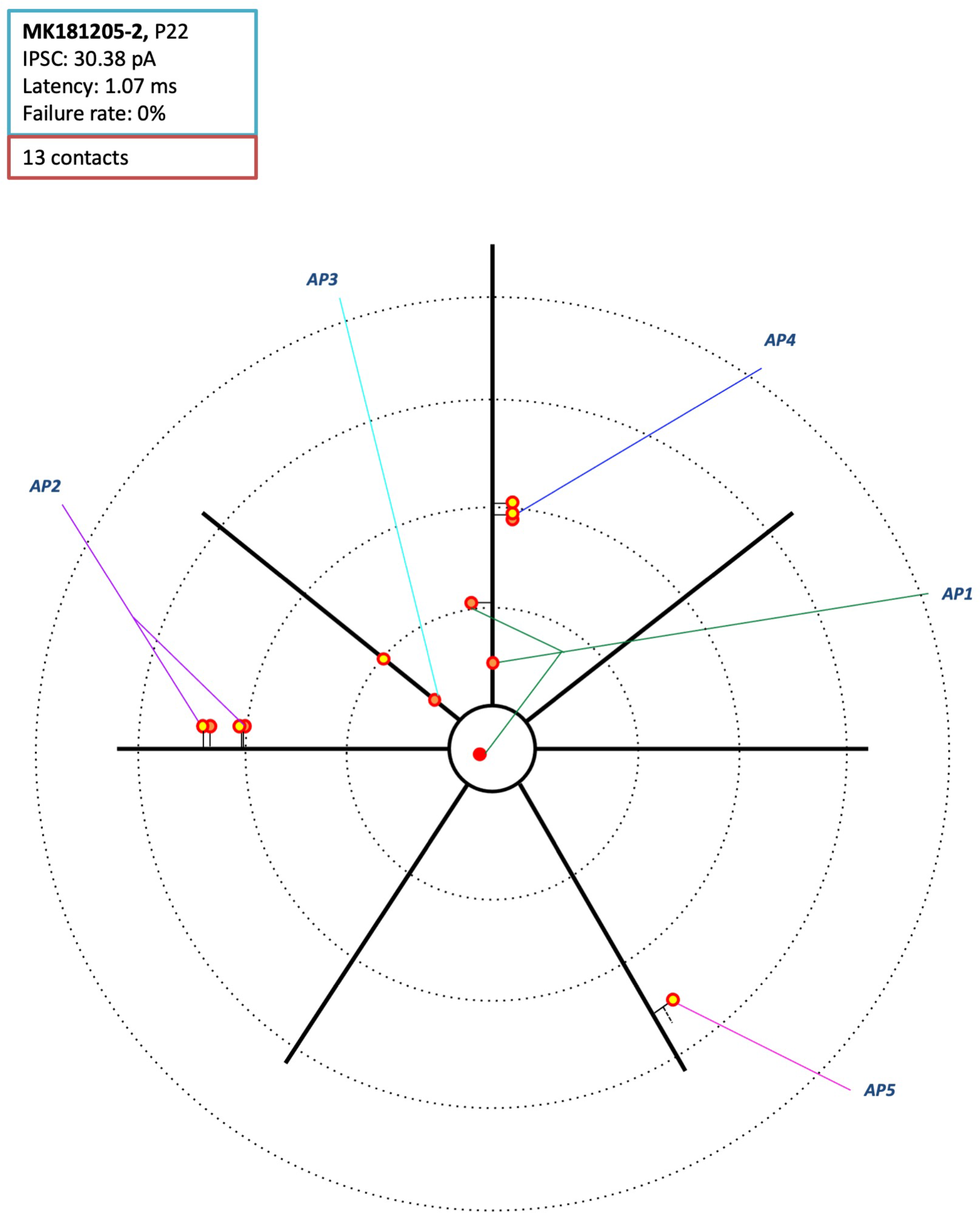
Axonograms of the presynaptic PV+ interneurons and dendrograms of the postsynaptic pyramidal neurons from the P14 and P22 neuronal pairs.

